# Heterogeneity in environmental stiffness alone can guide cells and shape tissues

**DOI:** 10.1101/2025.10.10.681597

**Authors:** Mathis Grelier, Carlos Ureña Martin, Maja Milas, Lital Bruskin-Pevzner, Yuval Segal, Madhura Ramani, Lovro Nuić, Guillaume Le Saux, Ana-Sunčana Smith, Mark Schvartzman

**Affiliations:** PULS Group, Department of Physics, Center for Computational Advanced Materials and Processes (CAMP), Friedrich-Alexander-Universität Erlangen-Nürnberg, Erlangen, Germany; Department of Materials Engineering and Ilse Katz Institute for Nanoscale Science and Technology, Ben-Gurion University of the Negev, Beer-Sheva, Israel; Group for Computational Life Sciences, Division of Physical Chemistry, Ruđer Bošković Institute, Zagreb, Croatia

**Keywords:** cell mechanics, stiffness patterns, contact guidance, mechanobiology

## Abstract

While topographical and chemical cues are well known to regulate cell shape and function, the role of stiffness heterogeneity has remained unclear. Here, we demonstrate – for the first time to our knowledge – that cells can be guided solely by the stiffness heterogeneity of their environment. To that end, we engineered a cell-guiding platform with abrupt, subcellular stiff and soft domains, whose flatness and uniform chemistry eliminated confounding cues. Cells elongate and align along stiff regions, sensing soft domains as barriers when wider than 2 microns. Perturbated myosin activity, cortical tension, and elasticity contrast reveal distinct biomechanical contributions, while a probabilistic model integrating adhesion, contractility, and cortical tension extracts key mechanical parameters characterizing the cellular state. Finally, experiments and dissipative particle dynamics demonstrate collective stiffness-based contact guidance. This work identifies stiffness heterogeneity as a fundamental regulator of cell and tissue organization and provides a framework for designing mechanoregulatory biomaterials.

## 1 Introduction

Shape, orientation, and motility of individual cells and tissues are vital for numerous biological processes [1]. Some examples are immune cell migration toward inflammatory sites, wound healing, embryogenesis, morphogenesis, and cancer metastasis [2]. Besides chemical signalling, these behaviors are guided by physical cues provided by the environment—foremost the elasticity, topography, and the spatial distribution of biochemical functionalities such as adhesion molecules or growth factors [3, 4]. When these cues form gradients, cells respond through haptotaxis [5, 6], durotaxis [7, 8, 9], or topotaxis [10]. When cues are spatially oriented without forming a gradient, cells align along them through contact guidance [11, 12]. For example, contact guidance underpins tendon nanofiber formation [13], cardiac muscle alignment for conduction and contraction [14], and dissemination of tumor cells [15]. It has also been exploited in tissue engineering—for dental implants [16], cochlear implants [17], and coronary stents [18]. Spatial heterogeneity in substrate stiffness and topography can, furthermore, induce directed cell migration on surfaces [19].

In natural environments, multiple cues often combine to produce collective guidance effects (Fig. 1a). For example, ECM deposition along topographical features aligns adhesive and topographical cues [20, 21, 22]. Oriented fibrillar ECM networks also induce contact guidance through mechanical anisotropy in a 3D matrix [12]. Chemistry can modulate material stiffness [23, 24], while the topography–stiffness crosstalk has been studied in significant detail [25] particularly in reconstituted systems. Ex vivo studies have, furthermore, proven valuable in dissecting individual mechanisms (Fig. 1b): durotaxis on elasticity gradients [26], haptotaxis on adhesive gradients [27], and topotaxis on topographical gradients [25]. Similarly, surfaces with ridges, grooves, or adhesive lines have demonstrated topographic or chemical contact guidance [28, 29, 30].

**Fig. 1.**
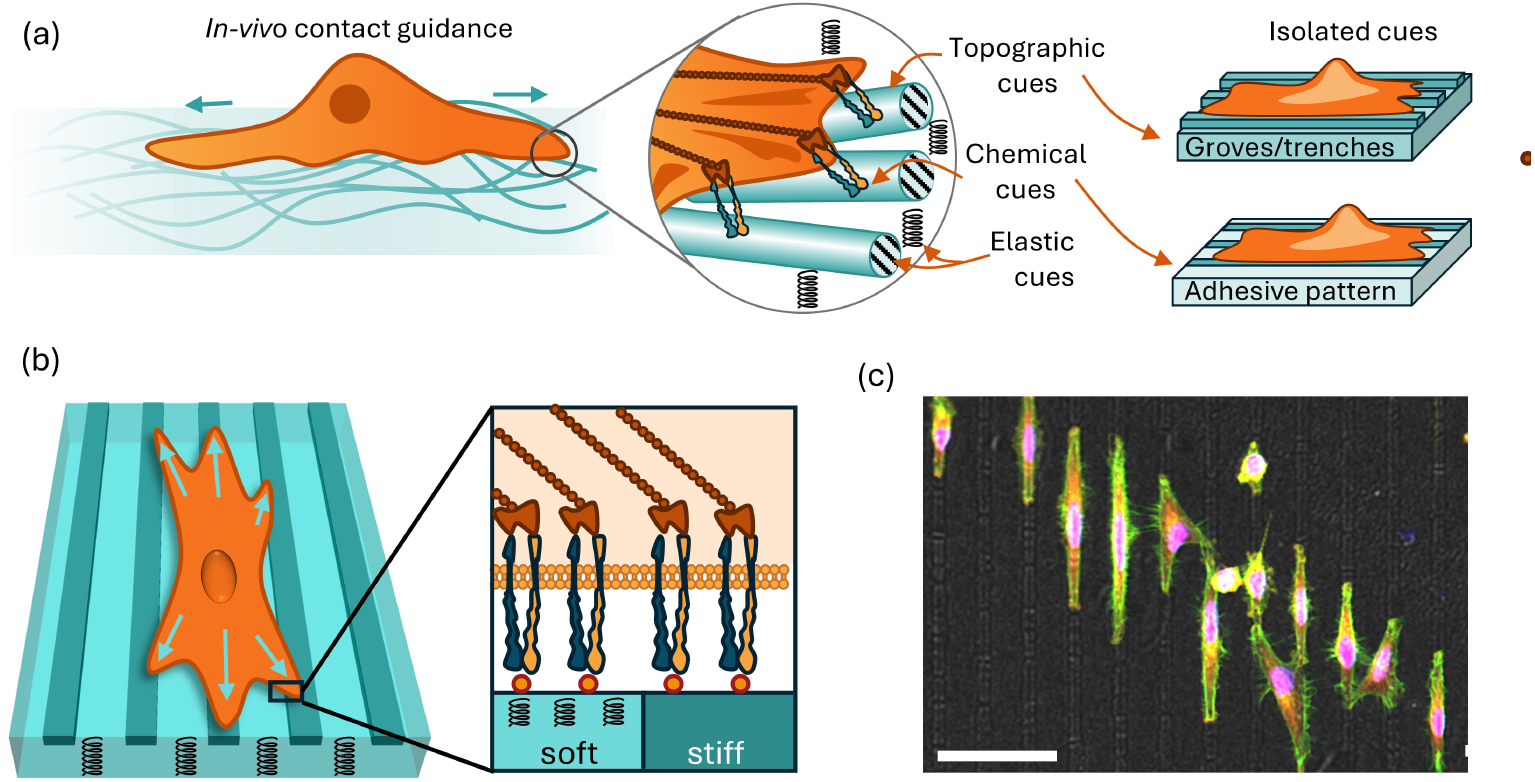
Shematic representation of contact guidance mechanisms. a) On the left we show contact guidance in natural environments relies on topological, adhesive and elastic variations (midle). Some of these mechanisms (topological and chemicaladhesive clues) were isolated in mimetic models and consequently studied in the literature b) Design of a model system to study contact guidance induced by contrast in elasticity, established herein. (c) Wide field image of cells deposited on a surface with vertical elasticity patterns clearly demonstrating that such design indeed induces contact guidance. Scale bar denotes 50 *µ*m. (d) Cell spreading on the elasticity pattern with stained nuclei (blue), actin (red) and paxilin (green). Scale bar denotes 10 *µ*m.

By contrast, contact guidance driven solely by substrate stiffness (Fig. 1b), where cells align in response to anisotropic elasticity without topographic or chemical cues, remains unexplored due to technical limitations. Specifically, an ex vivo system capable of isolating this effect would require a flat, chemically uniform surface patterned with sharply defined, subcellular-scale regions of varying stiffness. Such a configuration has not been achieved in previously reported cell-guiding substrates with spatial stiffness distributions [19, 31, 32, 33, 29], which, in particular, did not exclude topographies that could confound cell behavior. Consequently, a fundamental question remains: can stiffness heterogeneity alone guide cell alignment?

In this work, we show, for the first time, to the best of our knowledge, that substrate stiffness heterogeneity alone, without the involvement of other physical cues, can guide the shape and orientation of individual cells and cell agglomerates. To that end, we engineered flat, chemically uniform surfaces with alternating soft and stiff micro-stripes of subcellular width to isolate stiffness effects. Cells on these patterns aligned with the stripes, with the alignment strength increasing above a threshold stripe width (Fig. 1c,d). We rationalized these findings using a novel probabilistic theoretical approach suggesting that cells align with the stiffness patterns to minimize the energetic cost of crossing soft regions, balancing adhesion, tension, elasticity, and contractility. By softening stripes, as well as by inhibiting actin or myosin activity, we dissected the biomechanical contributions to this stiffness-driven contact guidance. Finally, using both agent-based simulations and model tissues, we show that elasticity patterns also direct collective cell behavior. This study provides new insights into cell mechanics and introduces a robust platform for ex vivo biomechanical research.

## 2 Cell-Guiding Stiffness Patterns

The primary requirement for detecting contact guidance driven purely by elasticity is the elimination of topographic features that could independently induce guidance. To meet this criterion, we fabricated flat elastic surfaces based on Polydimethysiloxane (PDMS) with embedded Silica stripes (Fig. 2a). The fabrication (described in detail in supporting information (SI)) included patterning of silica stripes on sacrificial silicon substrate coated with a thin Au film, using photolithography, silica evaporation, and lift-off. Then, thin film of liquid PDMS precursor, was cast over the pattern, cured, and cleanly detached from the silicon substrate by etching the gold layer, yielding soft PDMS surface with the bulk modulus around 1 MPa (unless stated otherwise), with embedded silica stripes of the bulk modulus around 50 GPa [34](Fig. 2b,c). Intentional 100 nm gaps were introduced along the silica lines to relieve mechanical stress in the PDMS and prevent the line cracking.

**Fig. 2.**
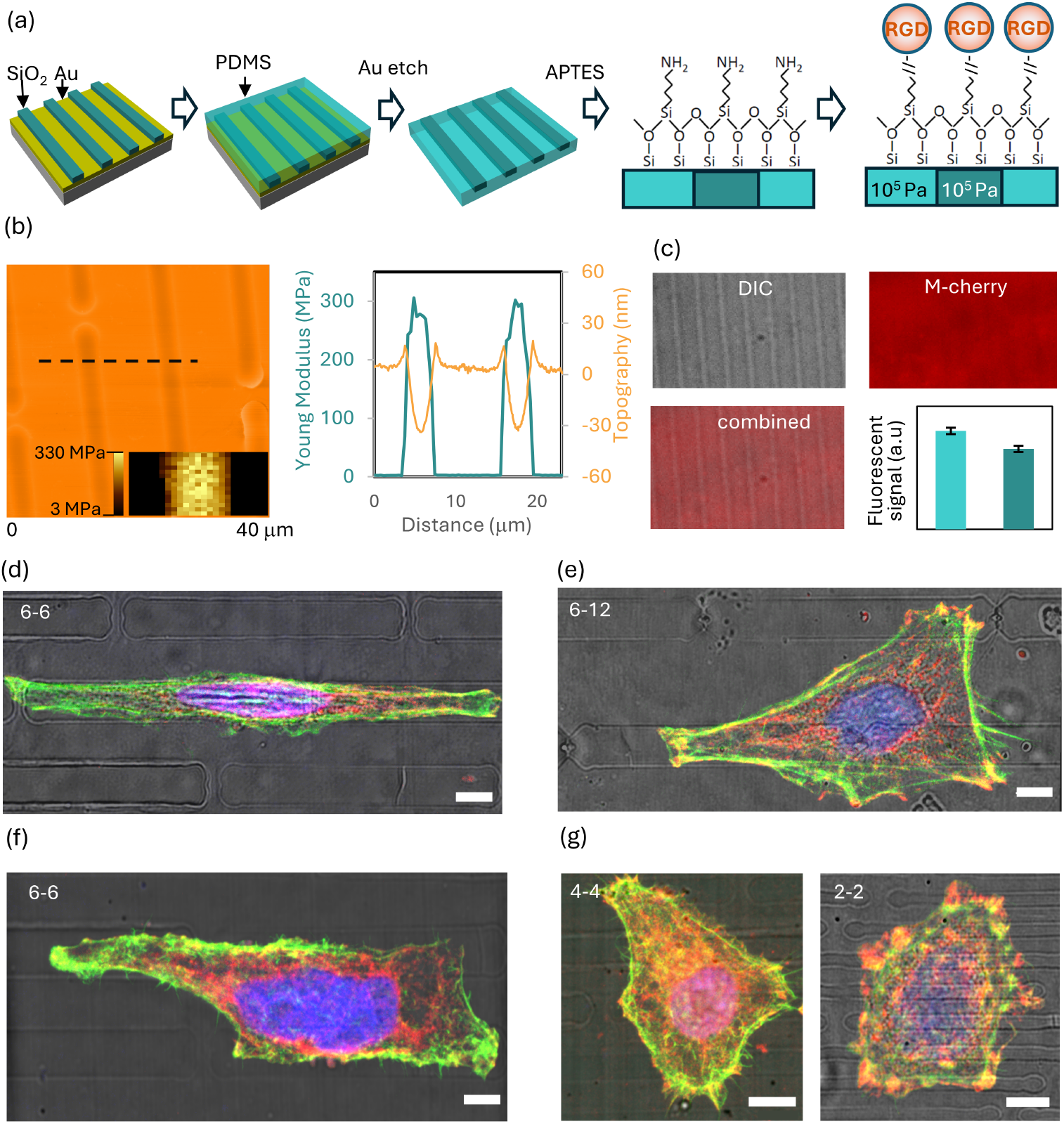
(a) Fabrication of flat and chemically uniform stiffness patterns. (b) AFM scanning of the stiffness pattern – 2D and profile. Inset in the 2 D scan shows elasticity mapping of a single stiff line and two neighboring soft lines. The plot on the right shows topography and stiffness profiles. (c) Functionalization uniformity: Differential Interference Contrast (DIC), fluorescent images, and their combinations are shown to visualize the uniformity of the chemisorption, quantitatively supported by the similarity of the fluorescent signal on the soft and stiffness. (d) A cell guided by the pattern that spread along single stiff lines, (e) - (f) Cells guided by the patterns bridging between neighboring stiff lines. (g) Cell lacking any guidance, with anisotropic spreading on narrow lines. Width of stiff lines (*L*_*h*_-*L*_*s*_) are shown in the left corner of each image. Scale bar denotes microns

Atomic force microscopy (AFM) was used to confirm the flatness of the resulting surfaces, with height differences between the soft PDMS and hard silica domains not exceeding 20 nm, which is well below the threshold known to induce topographic guidance at the line widths studied here [29]. Furthermore, the sharp contrast in elastic modulus across the pattern was characterized by AFM (Fig. 2b).

Overall, 12 distinct types of patterns were produced, featuring hard Silica line widths of *L*_*h*_ = 2, 4 or 6 microns, combined with soft PDMS lines of a width *L*_*s*_, with *L*_*s*_/*L*_*h*_ ratios of 1, 2, 3, and 4. Also, to verify that the observed 20 nm topography is indeed negligible in our experimental settings, and that cell guidance is solely attributable to stiffness variation, a set of control patterns with topographic lines of the same width and 20 nm height was fabricated entirely from either silica or PDMS (SI Section 1 and SI Fig. SI-1).

The second essential requirement, chemical uniformity, was achieved by chemisorbing (3-Aminopropyl) triethoxysilane (APTES) on both PDMS and silica domains, followed by conjugation with the RGD motif of fibronectin to facilitate cell adhesion. Uniform functionalization was confirmed by fluorescence microscopy using tagged molecules chemisorbed by the same process (Fig. 2a). These surfaces, therefore, provide an ideal model for studying contact guidance induced purely by elasticity patterns. Similar functionalization was applied to the Silica - and PDMS - based control surfaces with 20 nm topography.

These patterned and functionalized surfaces were used for overnight seeding of HeLa and NIH 3T3 fibroblast cells. The cells were then fixed, stained for nuclei, cytoskeleton, and vinculin, and imaged using microscope, about 100 cells of both cell types on each pattern. We, consequently, extracted the full distributions of morphological and orientational features *Q*_*n*_, such as cells’ area, perimeter, orientation relative to the stripe axis, and elongation for all patterns (approximately 1200 cells across 12 data sets) using Cellpose [35] and self-developed scripts (see GitHub and Zenodo archive associated with the manuscript).

Overall, cells showed clear elongation along the stripe direction, with distinct morphologies depending on pattern geometry. The two main spreading modes were: (i) cells confined almost entirely to a single stiff line, with adjacent soft lines restricting perpendicular spreading (Fig. 2d,f), and (ii) cells bridging across two or more stiff lines (Fig. 2e,g). In contrast, such morphologies were largely absent on patterns with narrow (2 *µ*m) lines, where cells spread mostly isotopically. Control surfaces with 30 nm topography showed no guidance effect, confirming that the observed behavior arises purely from elastic patterning.

## 3 Characterizing the mechanical state of the cells

### 3.1 Effective Steady State Energy Functional

To provide a model for this behavior, we rely on effective non-equilibrium steady state energy functionals, as an established tool to study cell mechanics (some classical examples include the vertex model [36, 37, 38, 39] or the cellular Potts model [40, 41, 42]). While the choice of the energy functional is not unique, in our framework, we focus on the simplest sufficient form, also discussed in previous work [43, 44]. Actually, more complex free energy functional forms have not proven to be more advantageous in our analysis. Accordingly, the cell’s energy functional *E*(***χ***), where the shape is parameterized by a vector ***χ*** is given by

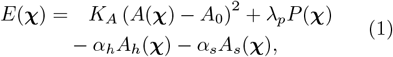

Here, *A* = *A*_*h*_ + *A*_*s*_ is the cell area and *A*_0_ its preferred projected area, with *K*_*A*_ denoting the elasticity modulus of the entire cell. This term captures the capacity of the cell to resist changes in its size, albeit in a simplified (2-dimensional) representation. The second term captures the contribution of cortical tension, where *P* is the perimeter and *λ*_*P*_ the associated line tension coefficient. It encodes the contractile forces from the actomyosin cortex and membrane tension projected into 2D, driving cells to optimize their perimeter for a given area. The last two terms encode the interaction between the cell and the substrate, comprising both adhesion, and adhesion-induced cytoskeleton contractility, with *α*_*h*_ and *α*_*s*_ characterizing the energetic preference (or cost) per unit area to create adhesive contacts with hard (*A*_*h*_) and soft (*A*_*s*_) regions of the substrate. The energy state **Ξ** of the cell ***χ*** is represented as **Ξ** = (*K*_*A*_ *λ*_*p*_ *A*_0_ *α*_*h*_ *α*_*s*_).

Consistently with the idea of an effective energy functional (1), we assume that the underlying statistics associated with fluctuations in cell shapes in the steady state *p*(***χ***) is Boltzmann-like

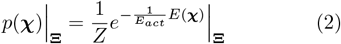

Here, *Z* is a normalization factor, integrating over all realization of cell shapes. Furthermore, we fix *E*_*act*_ = 1 to be the characteristic energy scale associated with cell activity in these systems, as in all experiments, the medium is the same, hence all cells are exposed to same nutrient levels (for a model with *E*_*act*_ being a free parameter, see SI Section 4).

To provide a versatile and compact basis that captures a broad spectrum of cell shapes ***χ***, we resort to the space of superellipses (Fig. 3a), parametrically defined as

**Fig. 3.**
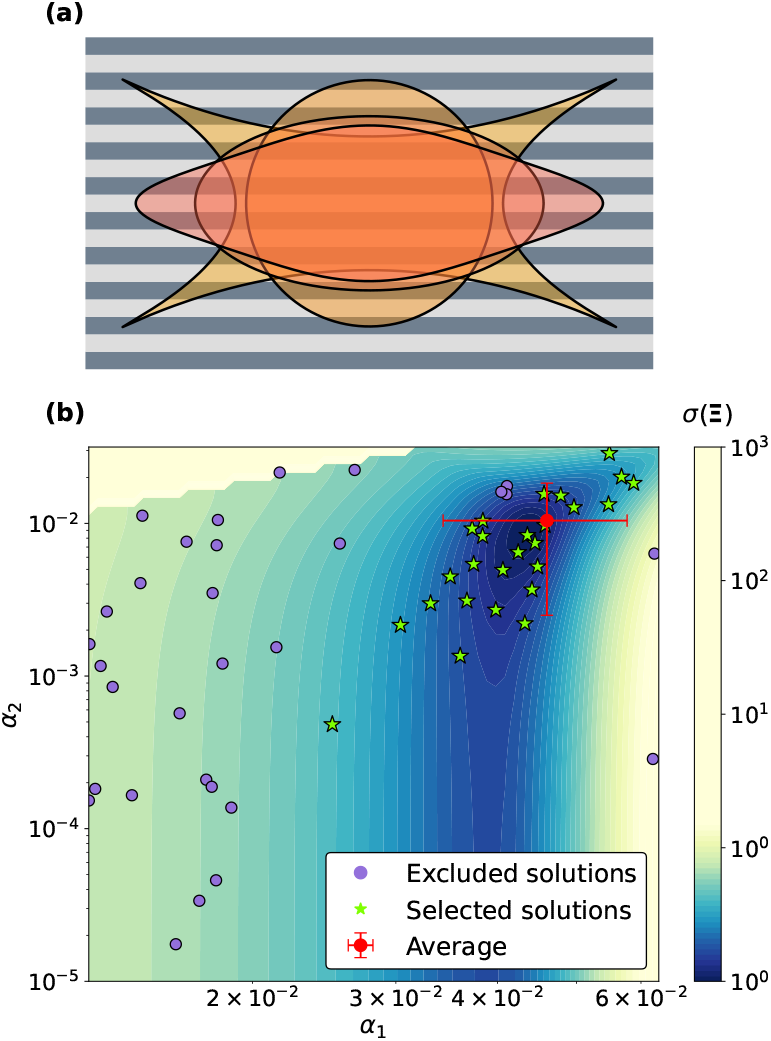
Optimisation of the parameters of the free energy functional. (a) Example of superellipses building the base for cell shapes, obtained for different value of *n* and *a* while maintaining a constant area (b) 2D map of the error function for the two parameters friction between the cell and the hard substrate *α*_*h*_ and the soft substrate *α*_*s*_, while the remaining parameters are kept constant at their average values, as used in Fig. 4. Dots represent the final results after the convergence of the optimization algorithm. The green stars indicate the optimal parameter values retained for calculating the average values of the parameters.

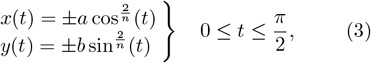

with *a, b* and *n* being parameters of the shape (see SI Section 2 for parametrization in the unisotropic settings of patterns).

This choice of a base balances biological reality with mathematical tractability, [45] enabling efficient exploration of the energetic landscape. Specifically, by setting **Ξ** to define *p*(***χ***), and integrating over all superellipses ***χ***, we can predict the average values of observables *Q*_*n*_

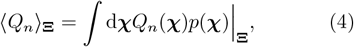

Consequently, we can provide model predictions for the average cell area *Q*_1_ = ⟨*A*⟩, aspect ratio *Q*_2_ = ⟨*c*⟩, and the distribution of cell orientations relative to the stripes *Q*_4_ = *P* (*θ*), along with the average orientation *Q*_3_ = ⟨*θ*⟩ (see SI Section 2).

### 3.2 Approach for determining the cell state parameters

With the effective energy specified, we use (perhaps developed?) a ***novel*** optimisation protocol to determine its parameters **Ξ** and capture theoretically the experimental observations. For this purpose, we employ the Globalized Bounded Nelder-Mead algorithm [46], which minimizes a preselected error function. As detailed in SI Section 3, we define this error function as the weighted quadratic difference between the model-predicted and experimental-associated averages of key cellular observables (area, orientation, and elongation), with weights given by the dataset’s size available for each observable (see eq. (S28)).

This approach is well-suited for multi-modal error landscape typical of cell mechanics, as it effectively navigates complex error surfaces containing multiple local minima. The globalized component systematically explores the parameter range as broad as several orders of magnitude by selecting not one but a sequence of initial parameter sets to increase the likelihood of escaping local minima and sampling unexplored regions of parameter space. Simultaneously, the bounded component ensures relevance and interpretability of the optimized parameters by constraining the search to physically meaningful ranges informed by values reported in the literature (Tab. 1) [43, 47, 48, 41, 49, 50, 51, 52].

To evaluate the sensitivity of the optimization protocol and the uncertainty of each free energy parameter comprised in **Ξ**, we repeat the optimization processes 100 times for each data set, building a set of {**Ξ}**. The average and standard deviations of the parameters are calculated from 30 optimisations that are clustered around the deepest minimum as shown in Fig. 3b. The obtained parameters are then used to generate the distributions of superelliptical cells, and calculate the set of *Q*_*n*_ shown jointly with measured distributions and experiments, as shown in Fig. 4. This ensures a robust and reliable comparison of our model with experimental data.

**Fig. 4.**
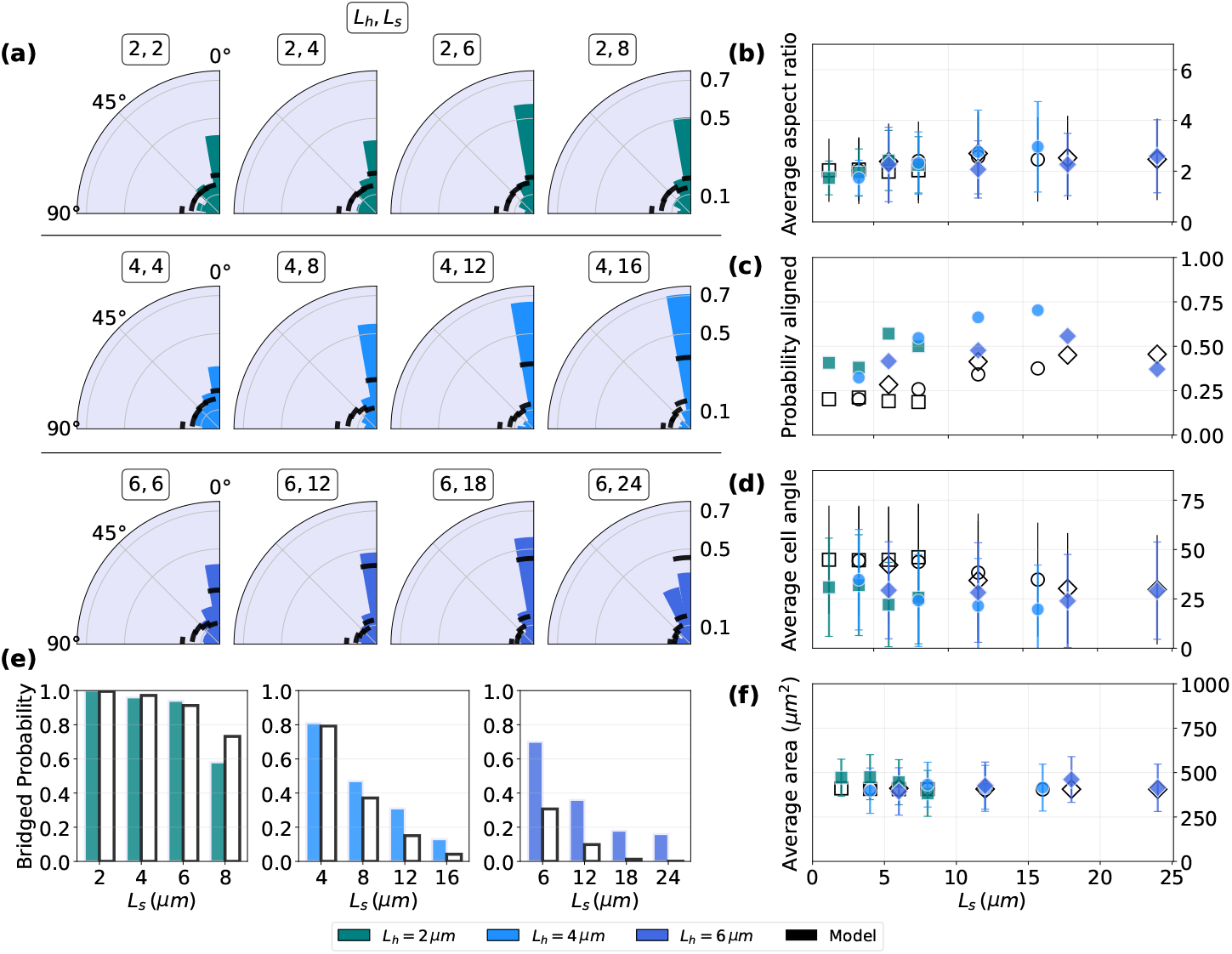
Characterizing the standard conditions. We show together experimental results (color-coded by *L*_*h*_) and model predictions (black lines and hollow symbols shaped as in experiments with the same *L*_*h*_) for varying widths of the soft substrate *L*_*s*_. (a) Alignment of cells relative to the substrate stripes orientation. (b) Average cell aspect ratio, reflecting changes in cell shape. (c) Alignment probability by evaluating the probability *P*_*θ<*15°_ to find a cell with a major axis closing less 15° with the pattern stripe. (d) Average orientation of the cells (e) Percentage of cells that extend across the soft substrate regions. (f) Average cell area as a function of the pattern. Experimental data in (a–d, f), shown by coloured symbols were used for parameter optimization (model result, hollow symbols), while (e) was measured independently (colored bars) and computed from the optimized model (hollow bars).

## 4 Cells on Elasticity Patterns

### 4.1 Cell morphology as a function of pattern geometry under standard conditions

With the experimental and theoretical methodologies established, we now examine how cells respond to patterns of different geometries, while simultaneously extracting all mechanical parameters that characterize them (Tab. 1), which is a key advantage of this analysis. Upon inspection of the results, we find that the cell maintains the same mechanical state even after adhering to different patterns, and that all components of {**Ξ}** spontaneously fall within the range expected from the literature. This means that cells utilise contact guidance to regulate their stiffness, contractility and adhesiveness.

**Table 1.** Ranges of the model parameters and corresponding fitted values.

| $\{\Xi\}$ | Unit | Range | Fitted value | Literature |
| --- | --- | --- | --- | --- |
| $K_A$ | N/m<sup>3</sup> | $10^7 - 10^{11}$ | $5.48 \times 10^8$ | [47, 51] |
| $\lambda_p$ | N | $10^{-9} - 10^{-8}$ | $9.81 \times 10^{-8}$ | [43, 41] |
| $A_0$ | $\mu\text{m}^2$ | $10^2 - 10^3$ | $3.87 \times 10^2$ | [50] |
| $\alpha_h$ | N/m | $10^{-3} - 10^{-1}$ | $4.78 \times 10^{-2}$ | [53, 47, 48] |
| $\alpha_s$ | N/m | $10^{-3} - 10^{-1}$ | $6.83 \times 10^{-3}$ | [53, 47, 48] |

In the context of efficiency of contact guidance by various patterns, we start our analysis from 2 *µ*m stiff stripe patterns (first row in Fig. 4a for HeLa cells, and fibroblasts in SI Fig. SI-5). Combined with 2 *µ*m soft stripes, both the experiments and the model show no cell alignment with the stripe direction. On these narrow soft and stiff lines, cells spread essentially isotropically, indicating that such patterns do not constrain cell orientation and fail to induce effective contact guidance. As the soft stripe width increases, experiments reveal a modest degree of alignment, which is slightly underestimated by the model (Fig. 4a,c,d).

As the width of the stiff stripes increases to *L*_*h*_ = 4 *µ*m and 6 *µ*m, both the experiments and the model quantitatively agree in showing a progressive increase of cell alignment with increasing soft stripe width, eventually reaching a plateau. This trend is particularly emphasized in the plot showing the fraction of cells oriented within 0*°* to 15*°* (Fig. 4c), and the plot of the mean orientation angle (where 0*°* denotes perfect alignment, Fig. 4d), corresponding to the most aligned segment in the wind rose charts (Fig. 4a). Alignment increases with soft stripe width, reaching saturation near 12 *µ*m, and remains largely independent of the stiff (silica) stripe width within the tested range.

This enhanced alignment (Fig. 4d) is accompanied by significant changes in cell shape. Specifically, cells elongate along the stiff substrate, while maintaining constant area in average (SI Fig. 4f). This modulation of shape reflects the cells’ tendency to maximize contact with the stiff parts of the pattern while being constrained by their own size. In this configuration, surface adhesion energy and cortical tension generated on the stiff stripes compensate for the lower contributions from the soft regions. Consistently, once the optimal *α*_*h*_ is reached, variations in *α*_*s*_ produce minimal changes to the error function optimizing the effective energy (Fig. 3b). This suggests that any pattern with sufficiently large soft stripes will promote alignment and elongation preferably along a single stiff line, unless crossing to the next stiff line is necessary to maintain cells compressibility and contractility.

### 4.2 Elasticity-based guidance is regulated by soft barriers

We further examine the capacity of soft stripes to act as energetic barriers in elasticity-based contact guidance by quantifying the percentage of cells that span two or more stiff silica stripes—i.e., form one or more bridges across soft regions (see Fig. 2e,f,g). We compare these experimental measurements with model predictions, by integrating systematically over all superelliptical cells configurations, weighed using the probability distributions parametrised by **Ξ**) The excellent agreement between the experiment and the model (Fig. 4e) underscores the predictive power of our framework, even for observables not included during parameter optimization.

In (*L*_*h*_, *L*_*s*_) = (2, 2) and (2, 4) patterns, the number of bridging cells is maximized, simply because the periodicity of the pattern of 4 and 6 *µ*m, respectively is less or similar to the typical length scale of the cell 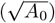. In this case the narrowness of the single soft stripe allows for the signal averaging by individual focal adhesions which cross both soft and hard backgrounds, and no contact guidance is observed (Fig. 2g). As the width of soft stripes increases, the signal associated with two elastic surfaces become better delineated. The number of bridging cells decreases showing that the soft stripes act as “spreading barriers” that broaden with the stripe width. Already for patterns (*L*_*h*_, *L*_*s*_) = (2, 8), the number of bridging cells is reduced, at which point cells also elongate appreciably along the narrow stiff stripe - i.e. to prevent bridging. Notably, this gives a clear length scale for lateral sensing of elastic properties of the surface by a single cell.

With stiff lines becoming wider, the barriers for bridging become deeper, as the signal to noise ratio of mechanosensing decreases. This is evident from the decrease in the number of bridging cells between patterns with increasing *L*_*h*_ at constant *L*_*s*_ (compare bridging probabilities of (*L*_*h*_, *L*_*s*_) = (2, 4) and (4, 4); (2, 6) and (6, 6); or (2, 8) and (4, 8)). When both effects combine, the stiff substrate being sufficiently wide to deepen the potential relative to the soft surface, and the width of the soft stripe being sufficient to extend the separation, the barrier becomes effectively impenetrable (*L*_*h*_, *L*_*s*_) = (4, 16) or (6, *L*_*s*_ *>* 12).

Together, these results demonstrate that substrate geometry directs cell orientation in a predictable, quantifiable manner. Notably, the absolute width of the soft strip, rather than the stiff/soft width ratio or stiff width alone, governs cell alignment, as captured by the effective energy approach. Consequently, using modeling, we can foresee the conditions under which contact guidance emerges or fails. This establishes a robust framework for mechanistically linking pattern geometry to collective cell orientation across a broad parameter space.

## 5 Actively modulating cell states

To further understand the regulation of stiffness-induced contact guidance and verify the consistency of our modeling approach, we assess how altering the balance of energetic contributions in our effective energy framework (eq. 1) modulates contact guidance relative to benchmark conditions. Experimentally, this is achieved by repeating the cell spreading assays for ensembles of HeLa cells under while evoking different mechanical conditions and measure the average cell orientation, aspect ratio, area, and probability of bridging in each case (Figs. SI-6, SI-7, SI-8). We then apply our optimization protocol to characterize the resulting states and predict cell shape distributions, enabling direct comparison with experiments and rationalizing the observed trends (Fig. 5).

**Fig. 5.**
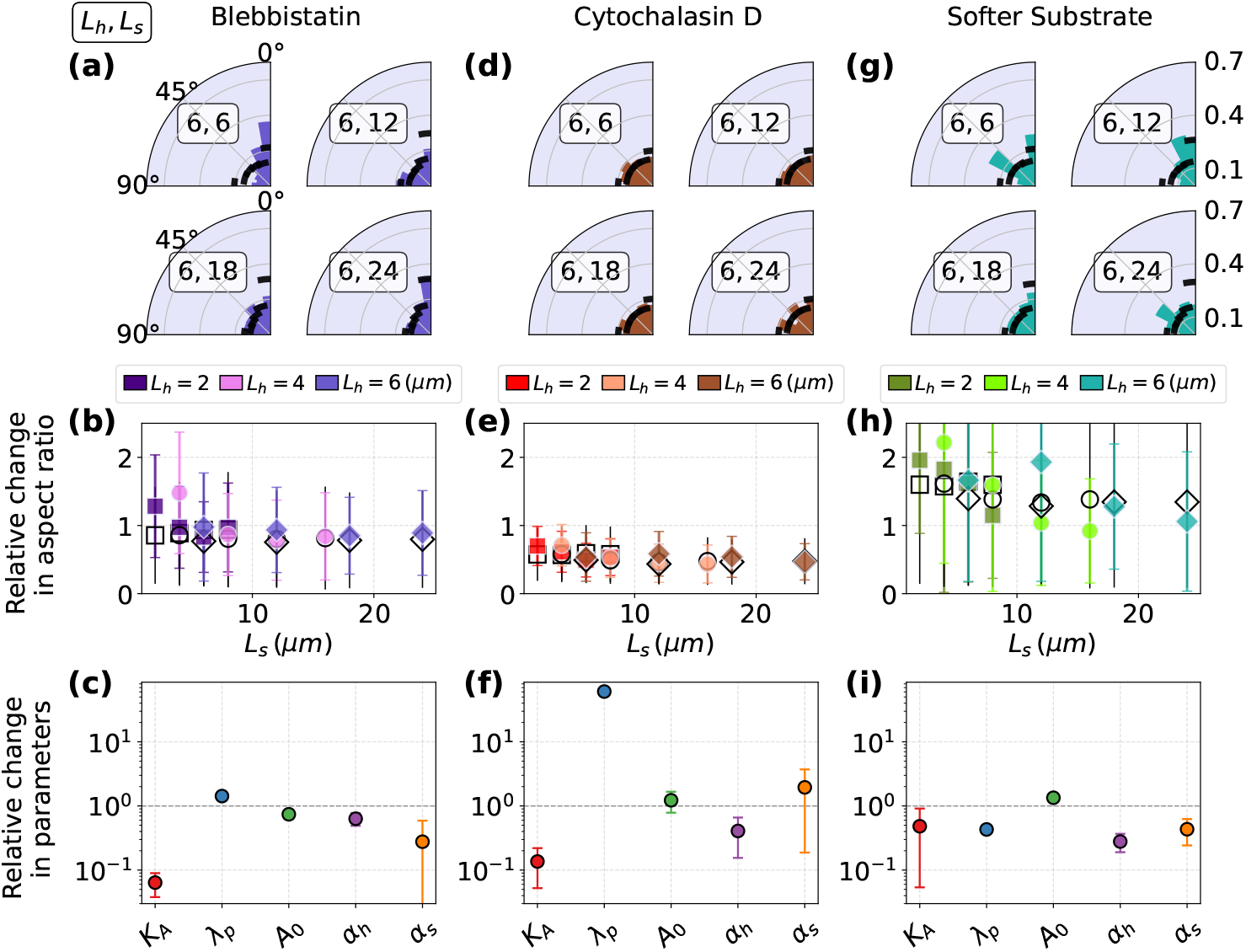
Comparison of treatments on cell orientation, elongation, and effective energy parameters. Columns correspond to the different perturbations (Blebbistatin, Cytochalasin D, Softer substrate), while rows represent successive levels of analysis. The first row shows cell orientation as a function of increasing soft stripe width *L*_*s*_ for a fixed hard stripe width *L*_*h*_ = 6, *µm*. The second row reports cell elongation for all couples (*L*_*h*_, *L*_*s*_), using the same markers and respective color code for each treatments as in Fig. 4. The third row presents the outcome of our free-energy optimization framework for each treatment. All quantities are normalized by the optimized parameters obtained from the reference experiments, enabling direct quantitative comparison and identifying which physical parameters of the cell are most affected by each treatment.

### 5.1 Affecting the capacity of a cell to deform and polarize

We first target the cell’s capacity to deviate from its preferred size (the first term in eq. 1) by treating cells with blebbistatin, [54, 55] a selective inhibitor of myosin II that reduces actomyosin contractility [56] (left panels in Fig. 5).

We find that on surfaces with elasticity patterns, blebbistatin treatment maintains cell alignment and spreading along stiff stripes with cells elongationg along the stripes as in standard conditions (Fig. 5a,b and Fig. SI-6b for non-normalized data). This finding aligns with previous reports showing that blebbistatin disrupts topographic contact guidance, [15, 57, 58], yet supports adhesive pattern-induced guidance [59, 60], potentially following the same mechanism.

This experimental observation is consistent with the finding obtained from our analysis of the effective steady state in which blebbistatin decreases the area elasticity modulus *K*_*A*_ while *λ*_*p*_ and adhesion parameters *α*_*h*_ and *α*_*s*_ remain unaffected (Fig. 5c. This suggest the disruption of actin support to the focal adhesion is compensated by membrane spreading and elongation along the stiffer substrate features.

### 5.2 Affecting effective tension

Cortical tension, represented by the second term in eq. 1, is modulated by treating cells with cytochalasin D, which disrupts actin filament polymerization, disorganizes the cytoskeletal network, and impairs the formation of stable protrusions necessary for polarization and directed migration. The result is a loss of stiffness-induced contact guidance in our experiments (see Fig. 5d), as reflected by a marked decrease in cell aspect ratio and alignment along stiff stripes (see Fig. 5e for data normalized to standard conditions and Fig. SI-7 for absolute results). This finding is consistent with previous reports on micropattern- and topography-based guidance. [61, 62]

Mechanistically, cytochalasin transforms the actin cytoskeleton into a more fluid-like structure, while increasing the bending stiffness of the membrane [56], such that the cell behaves as a relatively stiff fluid droplet. This is captured in our effective energetic model as a significant increase in the effective tension parameter (*λ*_*p*_), and a drop in *K*_*A*_ while adhesive parameters *α*_*h*_, *α*_*s*_ are remain unchanged (Fig. 5f). Despite global cytoskeletal softening, the consequence is a limited cell’s ability to maintain elongated shapes aligned with substrate cues and promoting a preference for more isotropic, circular shapes (Fig. SI-7).

### 5.3 Modulating adhesion-related energies

To investigate the role of adhesive terms in the effective energy (last term in eq. 1), we reduce the stiffness of the soft PDMS stripes by altering the resin-to-hardener ratio from 1:10 to 1:50, decreasing the bulk modulus of the soft stripes by approximately two orders of magnitude [63]. This change diminishes cell-substrate adhesion on the soft substrate [64], effectively strengthening the barrier presented by these stripes and decreasing but not fully disrupting contact guidance (Fig. SI-8 and Fig. 5g).

Mechanistically, within our effective free energy framework, reducing the stiffness of soft stripes favoring cell spreading over the stiff regions compared to the benchmark condition, particularly for patterns smaller than the characteristic cell size. This in turn increases contact guidance on narrow patterns relative to the standard conditions (Fig. 5h and Fig.SI-8), as the stronger contrast between soft and hard substrates generates a more robust mechanosensing signal. Conversely, on wider soft stripes, larger soft regions the impact is less significant since the position of the energy minimum is less sensitive to changes in *α*_*s*_ (Fig. 3b). As a result, the mechanical state of the cell is actually not significantly affected relative to the standard conditions, as confirming the insensitivity to *α*_*s*_ (Fig. 5i)

## 6 Collective effects in Stiffness Induced Contact Guidance

Motivated by evidence that collective cell migration [65] can arise from heterogeneity in topographical or adhesive cues [66, 67], as well as from gradients of topographic, adhesive, and elastic signals [68, 10], we explore whether stiffness patterns can similarly drive cooperative effects. We test this possibility by comparing agent-based simulations with experiments on tissues of Madin–Darby Canine Kidney (MDCK) epithelial cells grown on the same patterns as discussed for the single cell case. (For methodological details see SI Section 7.2).

MDCK tissues are a gold-standard epithelial model characterized by polarized monolayers with mature tight junctions, displaying collective migration behaviours closely resembling those of native epithelia [69]. To image these colonies over a extended periods of time, we developed a clone that stably expresses LifeAct-EGFP (SI section Y). These cells were seeded inside PDMS stencils (1.5 mm in diameter) on fibronectin-coated stiffness-striped substrates, allowed to reach confluence for 18 h, and then released by gently lifting the stencil. Time-lapse imaging was performed in a 37*°*C, 5% CO_2_ environmental chamber on a Nikon Ti-E inverted microscope fitted with a 10x objective. Phase-contrast and GFP fluorescence channels were captured every 1h for 40 hours.

Our simulations extend an established framework [70, 44], which we recently validated by quantitatively capturing the mechanosensitive growth dynamics of MDCK tissues on homogeneous glass and elastic substrates, where the colonies exhibited radial growth [71]. The difference between these two substrates is captured by two distinct friction coefficients: a high friction factor representing the glass background and a lower friction factor for the soft gels stripes. The stripe widths are set relative to the size (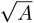 *≈* 4.6 *µ*m) of the largest cells observed in experiments, typically at the edge of the colony.

A most natural indicator of collective migration on striped patterns is the shape of a tissue as a whole. Starting from a circular colony, tissue elongation along a given axis serves as a reliable measure, since it indicates faster spreading in that direction. This effect can be very clearly seen in Fig. 6a where colonies growing on narrow stripes (*L*_*h*_ = *L*_*s*_ = 2 *µ*m) exhibit minimal changes in their aspect ratio, but colonies with stripes that show contact guidance on the single cell level, show preferential expansion along the stripes both in experiments and in simulations (Fig. 6b).

**Fig. 6.**
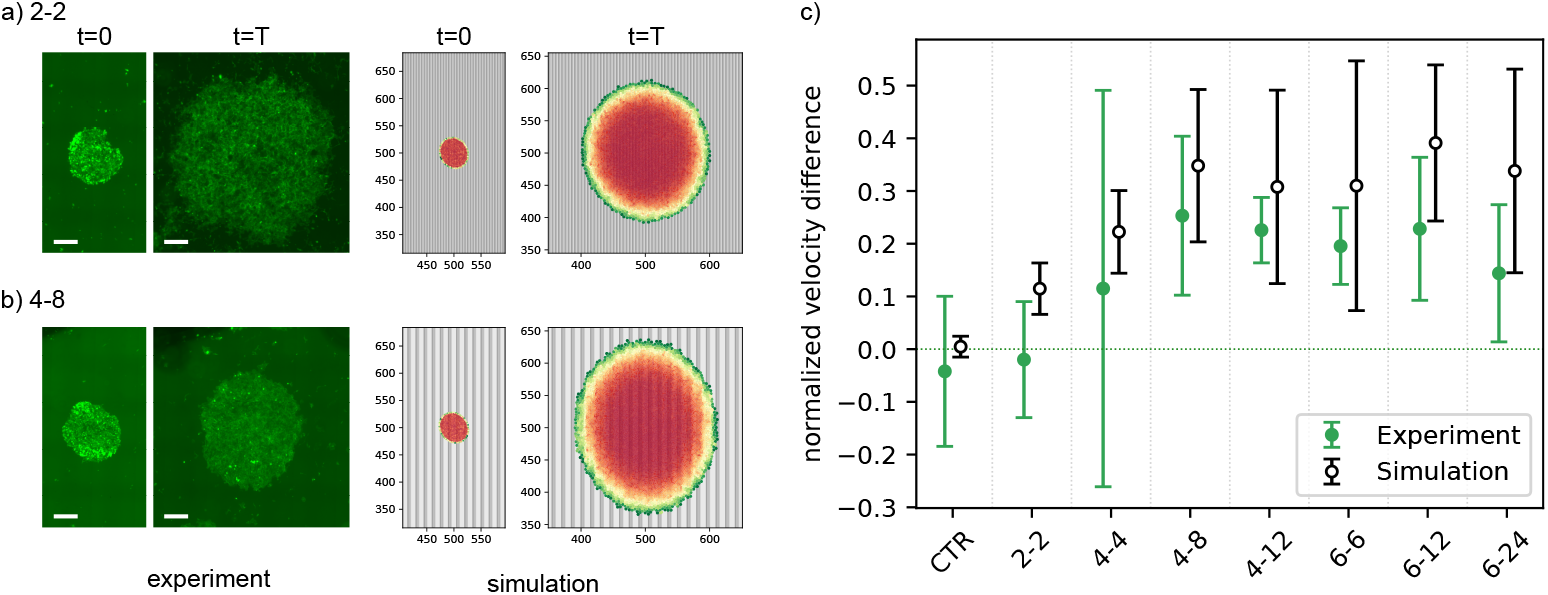
Contact guidance of tissues. (a,b) Snapshots of tissue growth on vertically oriented striped patterns (a narrow stripe pattern 2-2 and wider one 4-8). The first column displays experimental images processed using ImageJ (stripes not visible) while the second column shows corresponding snapshots from DPD simulations For each pair, the smaller image represents the initial and the larger one the final configuration. Simulations are initialized with an aspect ratio of 1.15, and the initial orientation angle of the tissue relative to the stripes is set to match the corresponding experimental condition. (c) The normalized velocity difference *ξ* quantifying the directional influence of the stripe pattern. The difference is positive for all simulated and most experimental patterns with sufficiently large stripes, indicating a promotion of vertical growth due to adhesion to stripes.

Due to small variations during preparations and removal of stencils, colonies of MDCK cells are often not perfectly circular, and the orientation of the major tissue axis is not necessarily aligned with the stripes at the beginning of the experiment. In this case, the aspect ratio of the colony is not a reliable measure of anisotropic collective motions, as it depends sensitively on time and the initial conditions. To circumvent this issue, we assess the difference in spreading velocity parallel and perpendicular to the stripes. As a relevant metric, we define the following anisotropic expansion parameter *ξ* = (*v* _‖_*−v*_*⊥*_)*/v*_0_, where *v*_⊥_ and *v*_‖_ are the velocity of the tissue front along the axis perpendicular and parallel with the stripes, respectively, while *v*_0_ denotes the velocity of the moving front of the tissue on a homogeneous substrate. With this definition, *ξ* = 0 denotes radial growth, while *ξ >* 0 indicates preferential expansion along the stripes.

In experiments and simulations, *ξ* is extracted from the data by first determining the confidence ellipse that encapsulates the tissue, measuring the temporal projections of changes in its principal axes (see SI Section 7.3 for technical details), and extracting *ξ* as a function of the pattern (Fig. 6c).

For narrow stripes smaller than the relevant cell size, we find that the tissue is insensitive to surface changes and the anisotropy of growth is weak. However, as soon as the stiff stripe reaches a width that is comparable to the characteristic cell size (*L*_*h*_ = 4 *µ*m), the extent of the anisotropy of growth saturates to positive values of *ξ*, and becomes insensitive to the pattern details. A closer examination of the simulation shows that substrate patterns do not alter the shape of the advancing density front, indicating that edge dynamics is primarily driven by the proliferation pressure (SI Fig. SI-12). In contrast, the dense tissue region develops non-uniformities due to different proliferation rates imposed by the stiffness of the substrate [71]. This effect leads to an accumulation of larger cells along the stripes, thereby determining collective elongation. The exception are very narrow stripes when the tissue averages signals over the pattern and responds to it as it would to a homogeneous surface, thus remaining circular [72].

## 7 Conclusion

Our work demonstrates that heterogeneity in substrate elasticity alone can direct cell and tissue organization. On chemically uniform, topographically flat surfaces patterned with alternating stiff and soft domains, cells preferentially elongate and align along stiff regions, while soft regions act as barriers until a critical width of 2 *µ*m is reached. This contact guidance is effective over pattern length scales up to 30 *µ*m, establishing a fundamental range for stiffness sensing. Perturbation experiments reveal that myosin inhibition and softening of the compliant domains enhance alignment and elongation along stiff regions, whereas disrupting cortical tension diminishes contact guidance. To mechanistically interpret these results, we developed a probabilistic model integrating adhesion, contractility, and cortical tension, which allows the simultaneous extraction of all major mechanical parameters characterizing the cellular mechanical state. As such, the variability of cells under certain conditions is treated as fluctuations around the determined steady state. At the tissue level, elasticity patterns bias collective migration and proliferation-driven growth, guiding overall tissue morphology according to the orientation of stiff domains. These findings establish stiffness heterogeneity as a central cue in cellular decision-making and introduce a robust platform for probing the mechanics of cell and tissue organization. Beyond fundamental insights, they provide a new conceptual and experimental basis for engineering microenvironments that control tissue architecture, with implications for regenerative medicine, cancer progression, and the design of mechano-responsive biomaterials.

## 8 Data availability

All data supporting the findings of this study are available in the Zenodo repository under accession number DOI:10.5281/zenodo.17306321. Utilized scripts are provided on GitHub under https://github.com/mathisg-ui/Heterogeneity-in-environmental-stiffness-alone-can-guide-cells-and-shape-tissues.git.

## 9 Acknowledgment

This work was partially funded by a joint project of the German (DFG) and French National Science Foundations 289/11-1. Further funding was provided by the DFG project SM 289/10-1 for collaboration with Israel, and the Israel Science Foundation, Project 2016/21. We thank P. Nowakowski for insightful discussions.

## 10 Author Contributions

MG established the theoretical model and analyzed the cell data. CUM performed all experiments, with the support of LB-P, YS, and GLS. MM performed tissues simulations, with the support of MG and adapting the code developed by LN. Tissue experiments were analyzed by MR and MG. ASS and MS conceived the project, and supervised the team.

## Notes

https://github.com/mathisg-ui/Heterogeneity-in-environmental-stiffness-alone-can-guide-cells-and-shape-tissues.git

### Competing Interest Statement

The authors have declared no competing interest.

