## Supplementary information for "Heterogeneity in environmental stiffness alone can guide cells and shape tissues"

**4 Active Energy Scale, $E_{\mathbf{act}}$** **13**

**5 Standard conditions for fibroblasts** **15**

**6 Modulating the state of cells** **16**

### 1 Experimental Methods

#### 1.1 Fabrication of the stiffness patterns

Silicon wafers were diced into $2.5 \times 2.5$ cm squares and cleaned by sonication in acetone for 5 min, followed by rinsing with ethanol. The cleaned substrates were sputter-coated with a 20 nm gold layer and subjected to photolithography using a negative photoresist, followed by pattern transfer through the sequential evaporation of $SiO_2$ (50 nm) and Ti (3 nm), and lift-off.

Liquid PDMS precursor was prepared from a Sylgard 184 base-to-curing agent mixture (Gelest) at a $10:1$ ratio (unless otherwise specified). The mixture was degassed, gently poured over the patterned silicon substrates placed in Petri dishes, and cured overnight in an oven at 60 °C. After curing, the PDMS was trimmed around the sample perimeter, and the PDMS–silicon sandwich was carefully removed from the Petri dish. The assembly was then immersed in a standard Au etchant ($KI/I_2$ solution) until the gold film was completely removed, allowing the PDMS with the embedded silica pattern to detach from the silicon.

#### 1.2 Functionalization of the pattern with RGD

PDMS–Silica patterns were first oxidized in Oxygen plasma for 30 s (Harrick PDC-32), and immediately immersed in a 5% ethanolic solution of (3-aminopropyl)triethoxysilane (APTES) for 30 min. The patterns were then rinsed thrice in neat ethanol, dried under a stream of nitrogen, and baked 30 min at 60 °C.

To functionalize the patterns with the Arg–Gly–Asp (RGD) peptide, samples were immersed for 48 h in an aqueous solution containing 0.2 M N-Ethyl-N′-(3-(dimethylamino)propyl)carbodiimide hydrochloride (EDC), 0.1 M N-Hydroxysuccinimide (NHS), 0.1 M 2-(N-morpholino)ethanesulfonic acid (MES) (EDC, NHS, MES from Merck), and 1 mM Gly–Arg–Gly–Asp–Ser (Genscript, USA). Samples were copiously rinsed with water, dried under a stream of nitrogen, and stored until further use.

Prior to cell experiments, the samples were sterilized by 30 min incubation in 70% ethanol, and after transfer to a sterile hood, copiously rinsed with sterile phosphate-buffered saline.

#### 1.3 Fabrication of control patterns

Two types of control samples were fabricated—one based on silica and the other on PDMS—both exhibiting the same $\sim$ 20 nm topography but lacking stiffness heterogeneity.

First, a $2.5 \times 2.5$ cm silicon wafer with a 200 nm thermal oxide layer was patterned by photolithography using the same method and mask described previously. The oxide layer was then gently etched through the mask in buffered oxide etchant (40% ammonium fluoride : 49% HF, $6:1$), diluted in water at a $1:5\, v/v$ ratio, to a depth of $\sim$ 30 nm. Removal of the photoresist with acetone revealed the first, rigid control pattern.

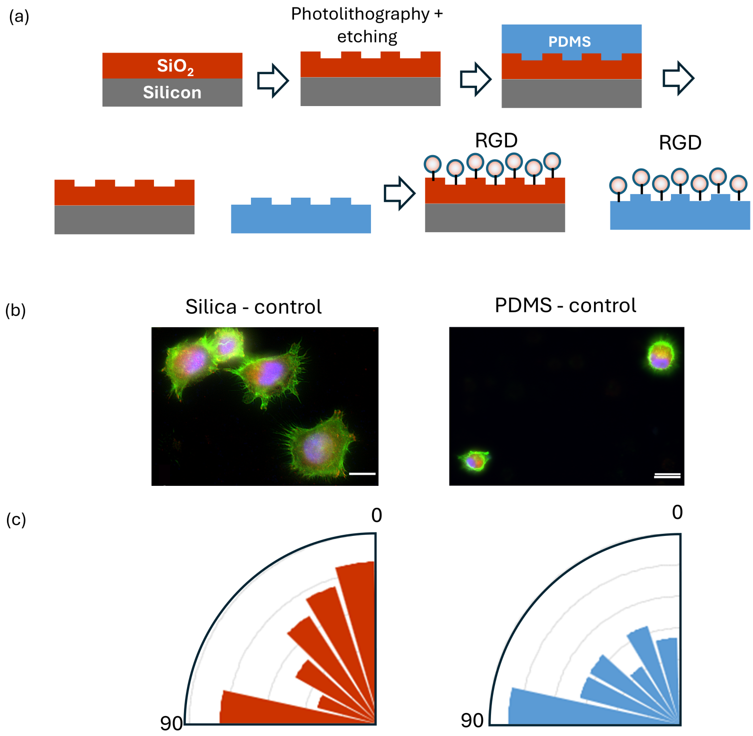

**Fig. SI-1** Control experiments show no contact guidance on samples with topography patterns of 20 nm. (a) Fabrication of the control samples. (b) Typical Hela cells after spreading on the control samples. Scale bars: 20 $\mu$m. (c) Distribution of the spreading direction on Silica and PDMS control samples.

This pattern was subsequently used to generate the second, soft control by replicating it in PDMS (Sylgard 184, Gelest) mixed with curing agent at a 10 : 1 ratio, cured overnight in an oven at 60 °C, and then peeled off. The Si/$SiO_2$ pattern was cleaned in piranha solution, and both patterns were functionalized with RGD as described above.

#### 1.4 Cell spreading on control patterns

HeLa cells were seeded onto the control surfaces and allowed to spread overnight following the same protocol used for the main experiments. The cells were then fixed, stained, and imaged using a fluorescence microscope. No significant alignment with the pattern was observed on either the stiff or soft control surfaces, confirming that topography at this level, in the absence of stiffness heterogeneity, is insufficient to induce cell contact guidance (Fig. SI-1b,c).

### 2 Basis of cell shapes and their morphological properties

#### 2.1 Parametrization of superellipses on the patterned surfaces

We define a superellipse in the $xy$-plane oriented at an angle $\theta$ relative to the stripes:

$$\begin{aligned} X &= x\cos\theta + y\sin\theta\,, \\ Y &= -x\sin\theta + y\cos\theta\,, \end{aligned} \tag{S1}$$

through the following parametric shape equation:

$$\begin{aligned} \frac{c+1}{4ac}\Bigg(&\big|(1-c)X + (1+c)Y\big|^n \\ &+ \big|(1+c)X + (1-c)Y\big|^n\Bigg) \le 1. \end{aligned} \tag{S2}$$

Here, the shape $\boldsymbol{\chi}$ is defined by four variables, $\boldsymbol{\chi} = (a, c, \theta, n)$. Specifically, $a = a'/(L_h + L_s)$ defines the normalized size of the cell along its principal axis ($a'$), perpendicular to the substrate stripes, relative to the characteristic length of the pattern ($L_h + L_s$). The parameter $c$ modulates the cell's elongation along the orthogonal axis, while $n$ controls the shape sharpness or degree of roundness: higher values of $n$ make the superellipse closer to a rectangle, whereas lower values yield more rounded contours.

#### 2.2 Morphological properties of superellipses

##### 2.2.1 Area

The area of such an object has an explicit solution and can be expressed in terms of the gamma function as:

$$A = 4ab\frac{\left(\Gamma\left(1+\frac{1}{n}\right)^2\right)}{\Gamma\left(1+\frac{2}{n}\right)}\,, \tag{S3}$$

with $a$ and $b$ the semi-diameters of the superellipse. To obtain shapes that more closely resemble the cells in the experiment, we set $a = b$ and apply a stretching transformation along the diagonal of the superellipse as follows:

$$S \cdot \begin{pmatrix} x(t) \\ y(t) \end{pmatrix} = \begin{pmatrix} c' & 1 \\ 1 & c' \end{pmatrix} \cdot \begin{pmatrix} x(t) \\ y(t) \end{pmatrix} = \begin{pmatrix} c'x(t) + y(t) \\ x(t) + c'y(t) \end{pmatrix} \equiv \begin{pmatrix} X(t) \\ Y(t) \end{pmatrix} \tag{S4}$$

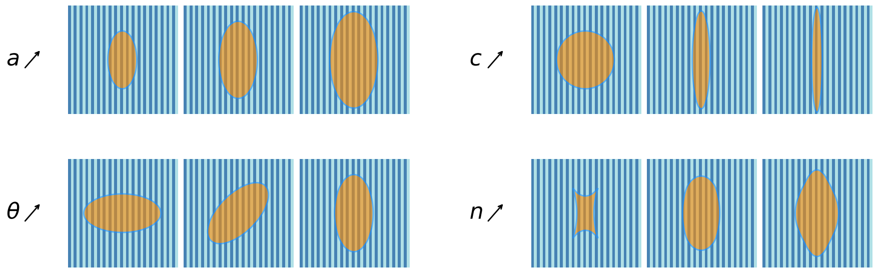

**Fig. SI-2** Shapes of the stretched superellipse defined in the main text with stripe widths $L_h = L_s = 4$ µm. Each variable increases along one row while the others are fixed at $a = 40.0$, $c = 2.0$, $\theta = \pi/4$, and $n = 2.0$.

where $c' = \frac{c-1}{c+1}$ is defined as a stretching coefficient ranging from 0 to 1, and $c$ is defined in the main text. The area of the stretched shape is then:

$$\begin{aligned} A_{\text{stretched}} &= |\det(S)|A \\ &= \left(c'^2 - 1\right) A \\ &= 4a^2 \left(c'^2 - 1\right) \frac{\left(\Gamma\left(1+\frac{1}{n}\right)^2\right)}{\Gamma\left(1+\frac{2}{n}\right)}. \end{aligned} \tag{S5}$$

The striped substrate can be divided into sets of hard ($S_h$) and soft ($S_s$) regions. However, computing which area of the superellipse belongs to which region is not trivial. Therefore, we use a sampling method by defining a grid of points and determining whether they are (i) inside the superellipse and (ii) located on $S_h$ or $S_s$. To do so, we use the Cartesian equation for a superellipse defined as

$$\left|\frac{X_c}{a}\right|^n + \left|\frac{Y_c}{b}\right|^n \leq 1, \tag{S6}$$

with $(X_c, Y_c)$ the coordinates of all points belonging to the rotated and stretched superellipse. After applying a rotation transformation (the superellipse is free to rotate by an angle $\theta$) and the stretching deformation defined above:

$$\begin{aligned} \begin{pmatrix} X_c \\ Y_c \end{pmatrix} = S^{-1}R\begin{pmatrix} x_c \\ y_c \end{pmatrix} &= \frac{1}{c'^2-1}\begin{pmatrix} c' & -1 \\ -1 & c' \end{pmatrix} \cdot \begin{pmatrix} \cos\theta & \sin\theta \\ -\sin\theta & \cos\theta \end{pmatrix}\begin{pmatrix} x_c \\ y_c \end{pmatrix} \\ &= \frac{1}{c'^2-1}\begin{pmatrix} \frac{c'(x_c\cos\theta + y_c\sin\theta) - (-x_c\sin\theta + y_c\cos\theta)}{(c'^2-1)a} \\ \frac{-(x_c\cos\theta + y_c\sin\theta) + c'(-x_c\sin\theta + y_c\cos\theta)}{(c'^2-1)a} \end{pmatrix} \end{aligned} \tag{S7}$$

where $(x_c, y_c)$ are the coordinates of all points belonging to the superellipse before any transformation. The search grid cannot be infinite in practice. Hence, we define a minimum rectangular search area centered at $(0, 0)$ (which can easily be shifted to

$(a_0, 0)$) with length $2l_a$ and width $2l_b$ for $n \geq 1$:

$$l_a = \Bigg[ \left| c' a \cos\left(z - \tfrac{\pi}{2}\right) + a \sin\left(z - \tfrac{\pi}{2}\right) \right|^{\frac{n}{n-1}} + \left| a \cos\left(z - \tfrac{\pi}{2}\right) + c' a \sin\left(z - \tfrac{\pi}{2}\right) \right|^{\frac{n}{n-1}} \Bigg]^{\frac{n-1}{n}} \tag{S8}$$

$$l_b = \Bigg[ \left| c' a \cos\left(z - \tfrac{\pi}{2}\right) - a \sin\left(z - \tfrac{\pi}{2}\right) \right|^{\frac{n}{n-1}} + \left| a \cos\left(z - \tfrac{\pi}{2}\right) - c' a \sin\left(z - \tfrac{\pi}{2}\right) \right|^{\frac{n}{n-1}} \Bigg]^{\frac{n-1}{n}} \tag{S9}$$

touching the borders of the superellipse. For $n < 1$, $n$ can be replaced by 1.01 to obtain a good approximation. For each of these points within this rectangle, we also check whether they belong to $S_h$ or $S_s$:

$$(x_c, y_c) \in S_h \Leftrightarrow \mod(x_c + a_0, L_h + L_s) \leq L_h \tag{S10}$$

$$(x_c, y_c) \in S_s \Leftrightarrow L_h \leq \mod(x_c + a_0, L_h + L_s) \;. \tag{S11}$$

The sampled areas can then be related to the exact stretched area (S5) as

$$A_h = \frac{N_h}{N_{\text{in}}} A_{\text{stretched}}, \;\; A_s = \frac{N_s}{N_{\text{in}}} A_{\text{stretched}}, \tag{S12}$$

with $N_h$ and $N_s$ the number of sampled points belonging to $S_h$ and $S_s$, respectively, and $N_{\text{in}}$ the total number of sampled points inside the superellipse.

##### 2.2.2 Perimeter

The computation of the energy potential requires the perimeter of the superellipse. As with the ellipse, no closed-form expression exists. Therefore, we formulate a parametric integral for the perimeter and approximate it numerically using Riemann sums. Superellipses have four sides defined by four parametric equations (3). Only two are necessary to compute the perimeter due to the symmetry of the shape. Let $P_1$ denote the length of the side defined by

$$\left. \begin{aligned} X_1(t) &= c' a \cos^{\frac{2}{n}}(t) + b \sin^{\frac{2}{n}}(t) \\ Y_1(t) &= a \cos^{\frac{2}{n}}(t) + c' b \sin^{\frac{2}{n}}(t) \end{aligned} \right\} \quad 0 \leq t \leq \frac{\pi}{2}. \tag{S13}$$

using (3) and (S4) with $x(t) = +a\cos^{\frac{2}{n}}(t)$ and $y(t) = +b\sin^{\frac{2}{n}}(t)$. The infinitesimal length of this curve is

$$dS = \sqrt{\left(\frac{dX_1}{dt}\right)^2 + \left(\frac{dY_1}{dt}\right)^2}\, dt\,. \tag{S14}$$

From this equation we obtain

$$\begin{aligned} P_1 &= \frac{2a}{n}\int_0^{\frac{\pi}{2}} \Bigg[\Big(\sin(t)^{\frac{2}{n}-1}\cos(t) - c'\cos(t)^{\frac{2}{n}-1}\sin(t)\Big)^2 \\ &\qquad + \Big(c'\sin(t)^{\frac{2}{n}-1}\cos(t) - \cos(t)^{\frac{2}{n}-1}\sin(t)\Big)^2\Bigg]^{1/2} dt \\ &= \frac{2a}{n}\int_0^{\frac{\pi}{2}} \Bigg[(1+c'^2)\Big(\cos(t)^{-2+\frac{4}{n}}\sin^2(t) + \cos^2(t)\sin(t)^{-2+\frac{4}{n}}\Big) \\ &\qquad - 4c'\cos(t)^{\frac{2}{n}}\sin(t)^{\frac{2}{n}}\Bigg]^{1/2} dt \end{aligned} \tag{S15}$$

Similarly, we define $P_2$ the other edge defined by:

$$\left.\begin{aligned} X_2(t) &= -c'a\cos^{\frac{2}{n}}(t) + b\sin^{\frac{2}{n}}(t) \\ Y_2(t) &= -a\cos^{\frac{2}{n}}(t) + c'b\sin^{\frac{2}{n}}(t)\,, \end{aligned}\right\} \quad 0 \le t \le \frac{\pi}{2}. \tag{S16}$$

and obtain:

$$\begin{aligned} P_2 &= \frac{2a}{n}\int_0^{\frac{\pi}{2}} \Bigg[(1+c'^2)\Big(\cos(t)^{-2+\frac{4}{n}}\sin^2(t) + \cos^2(t)\sin(t)^{-2+\frac{4}{n}}\Big) \\ &\qquad + 4c'\cos(t)^{\frac{2}{n}}\sin(t)^{\frac{2}{n}}\Bigg]^{1/2} dt \end{aligned} \tag{S17}$$

We identify $P_1 = P_-$ and $P_2 = P_+$ such that

$$\begin{aligned} P_\pm &= \frac{2a}{n}\int_0^{\frac{\pi}{2}} \Bigg[(1+c'^2)\Big(\cos(t)^{-2+\frac{4}{n}}\sin^2(t) + \cos^2(t)\sin(t)^{-2+\frac{4}{n}}\Big) \\ &\qquad \pm\, 4c'\cos(t)^{\frac{2}{n}}\sin(t)^{\frac{2}{n}}\Bigg]^{1/2} dt\,. \end{aligned} \tag{S18}$$

The total perimeter is then

$$\mathcal{P} = 2\,(P_- + P_+)\ . \tag{S19}$$

We approximate this integral by a Riemann sum as

$$\mathcal{P} = \int_0^{\frac{\pi}{2}} f(t)dt \approx \frac{\pi}{2N} \sum_{k=0}^{N} f\left(\frac{k\pi}{2N}\right) , \tag{S20}$$

with $N$ large enough to avoid large error.

##### 2.2.3 Elongation

For the elongation, we introduce the distance $d_1$ (respectively $d_2$) from the center to the middle of the edge defined by (S13) (respectively (S16)). Thus,

$$d_i = \sqrt{X_i\left(\frac{\pi}{4}\right)^2 + Y_i\left(\frac{\pi}{4}\right)^2} , \tag{S21}$$

where the factor $\frac{\pi}{4}$ comes from the fact that we want the distance to the middle of the respective edge defined between $0 \leq t \leq \frac{\pi}{2}$. We can show then that the elongation $r$ is

$$r = \frac{d_1}{d_2} = \frac{\sqrt{\left(c' a \cos\left(\frac{\pi}{4}\right)^{\frac{2}{n}} + b \sin\left(\frac{\pi}{4}\right)^{\frac{2}{n}}\right)^2 + \left(a \cos\left(\frac{\pi}{4}\right)^{\frac{2}{n}} + c' b \sin\left(\frac{\pi}{4}\right)^{\frac{2}{n}}\right)^2}}{\sqrt{\left(-c' a \cos\left(\frac{\pi}{4}\right)^{\frac{2}{n}} + b \sin\left(\frac{\pi}{4}\right)^{\frac{2}{n}}\right)^2 + \left(-a \cos\left(\frac{\pi}{4}\right)^{\frac{2}{n}} + c' b \sin\left(\frac{\pi}{4}\right)^{\frac{2}{n}}\right)^2}} \tag{S22}$$

which is simply:

$$r = \frac{1 + c'}{1 - c'} = c \tag{S23}$$

#### 2.3 Average quantities

As mentioned in the main, for any quantity $Q(\boldsymbol{\chi})$ we can compute its average value over all single cell shapes as

$$\langle Q \rangle = \sum_{\boldsymbol{\chi}} Q(\boldsymbol{\chi}) P(\boldsymbol{\chi}) . \tag{S24}$$

The average area $\langle A \rangle$ is computed as the sum of the area $A(a, c, n)$ defined by (S5), weighted by the probability distribution $P(a, c, \theta, n)$ over all possible shape configurations

$$\langle A \rangle = \sum_{a,c,\theta,n} A(a, c, n)\, P(a, c, \theta, n) . \tag{S25}$$

The average elongation $\langle c \rangle$ is given by

$$\langle c \rangle = \sum_{a,c,\theta,n} c\, P(a, c, \theta, n) . \tag{S26}$$

To obtain the probability distribution over $\theta$, or in other words the marginal probability distribution for the orientation, we sum over the other variables $a$, $c$ and $n$

$$P(\theta) = \sum_{a,c,n} P(a, c, \theta, n) \tag{S27}$$

These formulas (S25)–(S27) are used to generate the model plots shown in Fig. 4 of the main text, where we compare the derived quantities with the experimental data.

### 3 Optimization procedure

To quantitatively link our theoretical model to experimental data, we optimized the free energy parameter set $\boldsymbol{\Xi}$ by minimizing a tailored error function using the Nelder–Mead simplex method as implemented in the GNU Scientific Library (GSL). This algorithm belongs to the class of gradient-free minimizers and relies only on function evaluations at successive parameter vectors.

#### 3.1 Optimization algorithm

The Nelder–Mead method constructs an $n$-dimensional simplex from the initial guess $\mathbf{x}_0$ and step-size vector $\mathbf{s}$. At each iteration, the simplex adapts through geometrical transformations (reflection, expansion, contraction, or multiple contraction) applied to the vertex associated with the largest function value. This iterative process guides the simplex toward the minimum, while its characteristic size shrinks as convergence is approached. We employed the `gsl_multimin_fminimizer_nmsimplex2` routine from the GSL library.

#### 3.2 Error function

The discrepancy between model and experiment was quantified by a weighted quadratic error function,

$$\begin{aligned}\sigma(\boldsymbol{\Xi}) = w_A \left(\frac{\langle A_{\mathrm{model}}\rangle - \langle A_{\mathrm{exp}}\rangle}{A_{\mathrm{exp0}}}\right)^2 + w_c \left(\frac{\langle c_{\mathrm{model}}\rangle - \langle c_{\mathrm{exp}}\rangle}{c_{\mathrm{exp0}}}\right)^2 \\ + w_{P_\theta} \sum_\theta \Big(P_{\mathrm{model}}(\theta) - P_{\mathrm{exp}}(\theta)\Big)^2\end{aligned} \tag{S28}$$

where $w_A$, $w_c$ and $w_{P_\theta}$ are weights that are inversely proportional to the number of data points for each quantity. The dependence $\boldsymbol{\Xi}$ is encoded in $\langle A_{\mathrm{model}}\rangle$, $\langle c_{\mathrm{model}}\rangle$ and $P_{\mathrm{model}}(\theta)$ and has been omitted for the sake of clarity. Additionally, each term is normalized by the characteristic order of magnitude of its respective quantity, ensuring that all quantities contribute equally to the optimization. The goal of this error function is two-fold. On the one hand, it allows us to see the impact of the physical parameters grouped in $\boldsymbol{\Xi}$ on the difference between the model and the experimental results. On the other hand, the optimization determines interpretable and comparable value for these parameters.

#### 3.3 Domain constraints

To ensure physically meaningful parameters, the optimization was performed under box constraints implemented via the *extreme barrier* method [1]. In this approach, parameter vectors outside the admissible domain $D$ are assigned an infinite penalty,

$$\sigma_E(\boldsymbol{\Xi}) = \begin{cases} \sigma(\boldsymbol{\Xi}) & \text{if } \boldsymbol{\Xi} \in D, \\ +\infty & \text{otherwise,} \end{cases} \tag{S29}$$

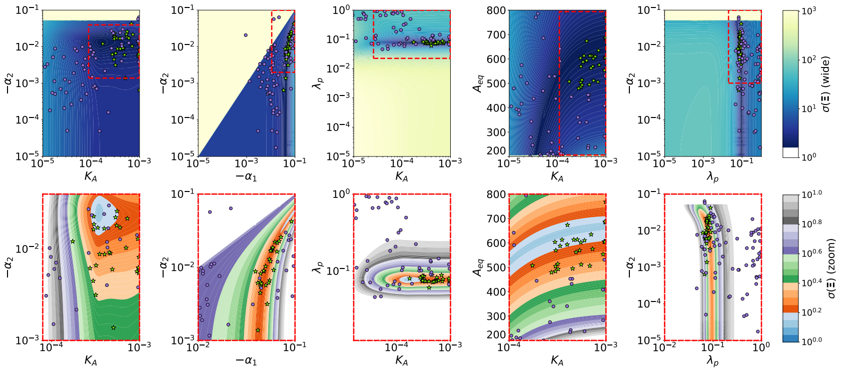

**Fig. SI-3** Two-dimensional parameter maps showing the dependence of the error function $\sigma(\Xi)$ on selected pairs of parameters. Each column corresponds to one parameter pair, with the top row showing the full parameter range (wide view) and the bottom row showing the corresponding restricted parameter domain (zoom view, indicated by the red dashed rectangle). Colors indicate the value of $\sigma(\Xi)$ on a logarithmic scale. Green star symbols denote the best $N = 30$ parameter sets with the lowest $\sigma(\Xi)$, while purple dots denote other candidate parameter sets after optimization. The zoom panels highlight the region of parameter space where optimal solutions concentrate.

with

$$D = \left\{ \Xi \subset \mathbb{R}^n \,\middle|\, K_A \geq 0,\ \lambda_P \geq 0,\ A_{\text{eq}} > 0,\ \mathcal{P}_{\text{eq}} > 0,\ \alpha_1 > 0,\ \alpha_2 > 0,\ \alpha_1 > \alpha_2 \right\}. \tag{S30}$$

as well as for the $\Xi$ outside of the range mentioned in Tab.1 in the main text.

#### 3.4 Robustness analysis

To assess robustness and parameter uncertainty, the optimization procedure was repeated 100 times for each dataset with different randomized initializations. The resulting parameter sets $\Xi$ were then studied, and statistical analysis (average and standard deviation) was restricted to the 30 realizations located in the deepest minimum (see Fig. SI-3). This procedure provides a reliable estimate of parameter variability while filtering out spurious local minima.

To further examine the structure of the parameter space, we mapped the objective function $\sigma(\Xi)$ along selected two-dimensional parameter planes, fixing all remaining parameters to their average values within the best solutions. As shown in Fig. SI-3, this reveals well-defined basins of attraction around the optimal region, indicating that the optimization landscape is not only sharp but also consistent across multiple parameter combinations. These maps confirm that the optimal solutions are localized and robust against small perturbations in parameter values.To gain a clearer view of the optimal region, we also present zoomed versions of the 2D maps, where the color scale is restricted to the range $[1, 10]$, and values exceeding 10 are shown in white. Small discrepancies arise between the optimal points and the minima displayed in the 2D maps, due to the fact that the maps are generated by fixing all other parameters to their average values. These average values differ slightly from the specific parameter sets at which each optimization run converged.

#### 4 Active Energy Scale, $E_{\text{act}}$

In the main text, we employ an effective energy functional, $E(\boldsymbol{\chi})$, to model cell mechanics and assume the underlying statistics of cell shapes, $P(\boldsymbol{\chi})$, follow a Boltzmann-like distribution as shown in Eq. 2. This approach is well-established for describing active biological systems that are inherently out of thermal equilibrium [2, 3, 4]. The parameter $E_{\text{act}}$ in the Boltzmann factor, analogous to an effective temperature, encapsulates the energy scale of all non-thermal, active processes within the cell, such as cytoskeletal remodeling and motor protein activity. It sets the characteristic energy of cellular fluctuations.
In our primary analysis, we fixed this active energy scale to $E_{\text{act}} = 1$, thereby defining it as our reference energy unit. Once fixed, it reduces the dimensionality of the parameter space and helps to avoid overfitting, allowing for a more robust determination of the other physical parameters in the energy functional

$$\Xi = (K_A,\ \lambda_p,\ A_0,\ \alpha_h,\ \alpha_s)\ .$$

To rigorously justify this simplification *a posteriori*, we conducted a supplementary, comprehensive optimization study where $E_{\text{act}}$ was included as a sixth free parameter alongside the components of $\Xi$. We then compared the results of this extended optimization with the results obtained in the main text where $E_{\text{act}}$ was fixed.
First, we assessed the impact of treating $E_{\text{act}}$ as a free parameter on the converged values of the other energy functional parameters. We ran the full optimization (with all six parameters free) 1000 times—ten times more than in the main analysis—to account for the enlarged parameter space. We then took the best 30 converged parameter sets $\{\Xi_{\text{free}}, E_{\text{act,free}}\}$ and compared the values of $\Xi_{\text{free}}$ to the corresponding values $\{\Xi_{\text{opt}}\}$ obtained from the original optimization.
Figure SI-4a shows the ratio of each parameter in $\Xi_{\text{free}}$ to its counterpart in $\Xi_{\text{opt}}$. The value for all five parameters are close to 1, with the bigger difference for $\alpha_s$ due to a larger standard deviation. This indicates that the values of the elasticity modulus ($K_A$), surface tension ($\lambda_p$), preferred area ($A_0$), and adhesion preferences ($\alpha_h, \alpha_s$) are remarkably insensitive to whether $E_{\text{act}}$ is fixed or allowed to vary. The minimal deviation observed demonstrates the robustness of our original parameterization and confirms that fixing the energy scale does not introduce a significant bias into the determination of the cell's physical properties.
Next, we analyzed the distribution of the converged values for $E_{\text{act}}$ itself from the full optimization. Figure SI-4b displays the probability distribution of $E_{\text{act,free}}$ for the 30 best-performing parameter sets. The distribution lies between 0 and 2, remaining within the unity range. This result is significant because it shows that, even without any prior constraint, the optimization process consistently identifies a characteristic active energy scale inherent to the cell type and experimental conditions.
Finally, to further isolate and confirm the value of the active energy scale, we performed a third set of optimizations. In this case, we fixed the parameters of the energy functional to their optimal values found in the main text, $\Xi_{\text{opt}}$, and treated only $E_{\text{act}}$ as a free parameter. This procedure effectively asks: given the optimal description

of the system's energy landscape, what is the most probable magnitude of cellular fluctuations?
The resulting distribution of optimized $E_{\text{act}}$ values is shown in Fig. SI-4c. The distribution converge to a single point, yielding the optimal value of $E_{\text{act}}$ for the previously determined optimal value of $\Xi$. This demonstrates that when the cell's mechanical and adhesive properties are correctly specified, the level of activity required to explain the experimentally observed shape distributions is consistently and precisely determined to be $E_{\text{act}} \approx 0.98$.

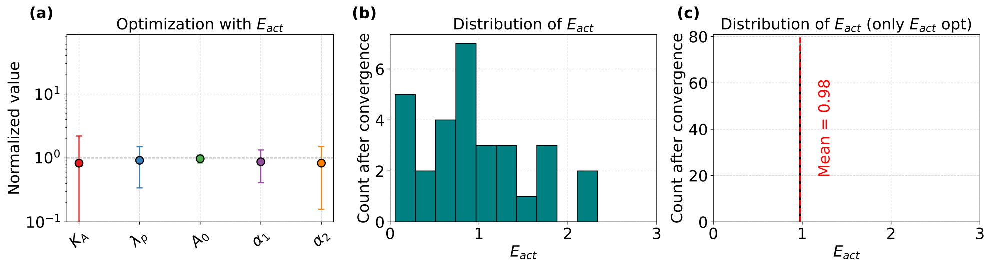

**Fig. SI-4** (**a**) Distribution of the ratio of parameter values obtained from an optimization where $E_{\text{act}}$ was a free parameter ($\boldsymbol{\Xi}_{\text{free}}$) versus the values from the main analysis where $E_{\text{act}}$ was fixed ($\boldsymbol{\Xi}_{\text{opt}}$). (**b**) Distribution of the optimized $E_{\text{act}}$ values from the full, unconstrained 6-parameter optimization. (**c**) Distribution of the optimized $E_{\text{act}}$ values when the other five parameters were held constant at their previously determined optimal values.

### 5 Standard conditions for fibroblasts

As for the control sample shown in Fig. 3 of the main text, we use the same procedure to fit experiments for different conditions with our model.

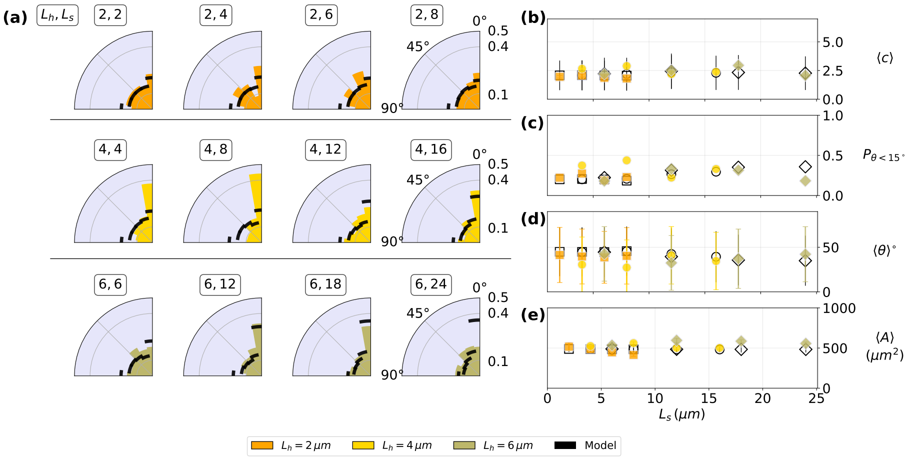

**Fig. SI-5** Fibroblasts on stiffness patterns under standard conditions. Comparison between experimental results (color coded by $L_h$) and model predictions (black, and hollow symbols) for varying widths of the soft substrate $L_s$. (a) Alignment of cells relative to the substrate stripes orientation . (b) Average cell elongation, reflecting changes in cell shape. (c) Percentage of aligned cells (d) Average cell orientation. (e) Probability for bridging between stiff lines. (f) Average cell area.

### 6 Modulating the state of cells

#### 6.1 Treatment with blebbistatin

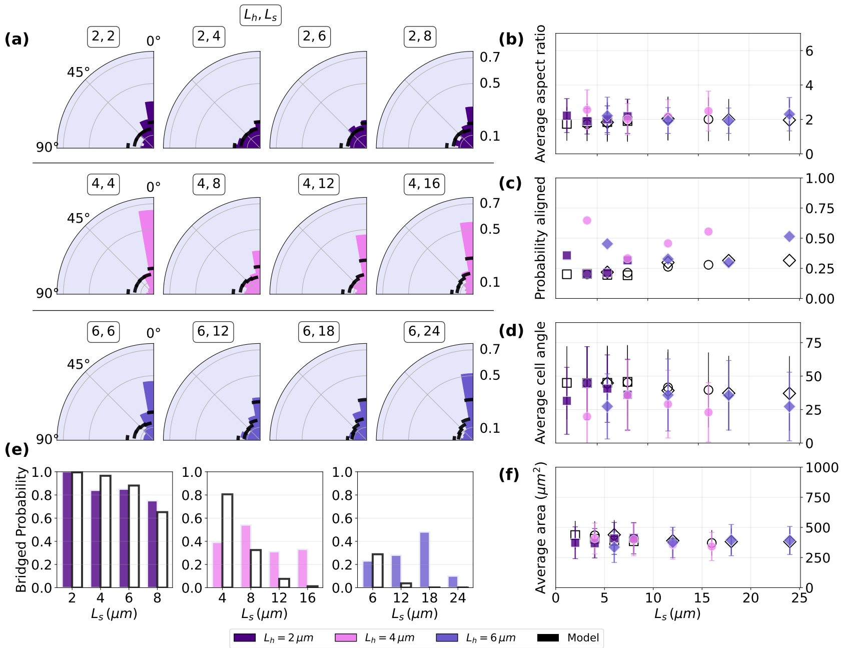

**Fig. SI-6** Contact guidance of HeLa cells treated with blebbistatin to make cells more flexible. Comparison between experimental results (color coded by $L_h$) and model predictions (black, and hollow symbols) for varying widths of the soft substrate $L_s$. (a) Alignment of cells relative to the substrate stripes orientation . (b) Average cell elongation, reflecting changes in cell shape. (c) Percentage of aligned cells (d) Average cell orientation. (e) Probability for bridging between stiff lines. (f) Average cell area.

#### 6.2 Treatment with cytochalasin A

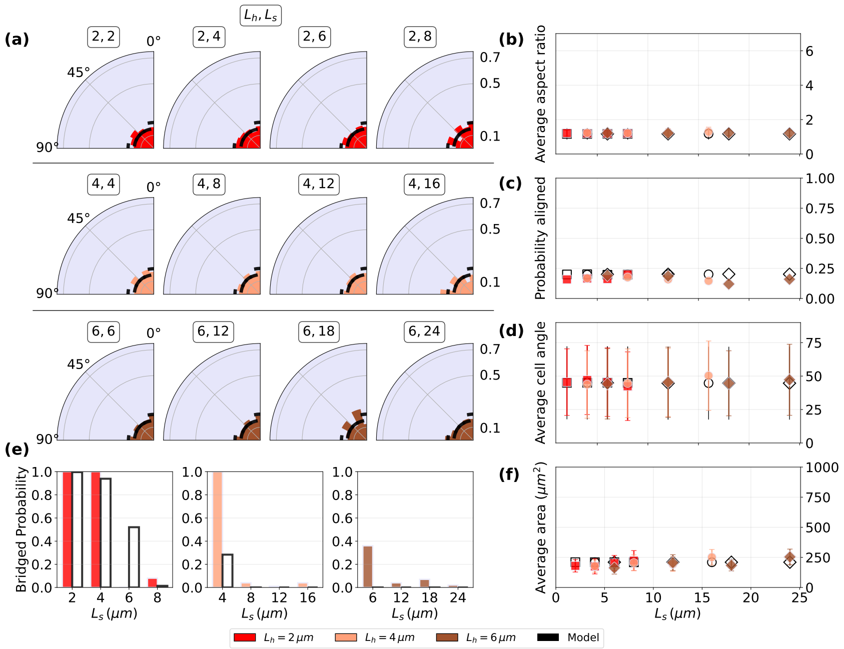

**Fig. SI-7** Contact guidance of HeLa cells treated with cytochalasin D to make cells less tense. Comparison between experimental results (color coded by $L_h$) and model predictions (black, and hollow symbols) for varying widths of the soft substrate $L_s$. (a) Alignment of cells relative to the substrate stripes orientation . (b) Average cell elongation, reflecting changes in cell shape. (c) Percentage of aligned cells (d) Average cell orientation. (e) Probability for bridging between stiff lines. (f) Average cell area.

#### 6.3 Making soft stripes softer

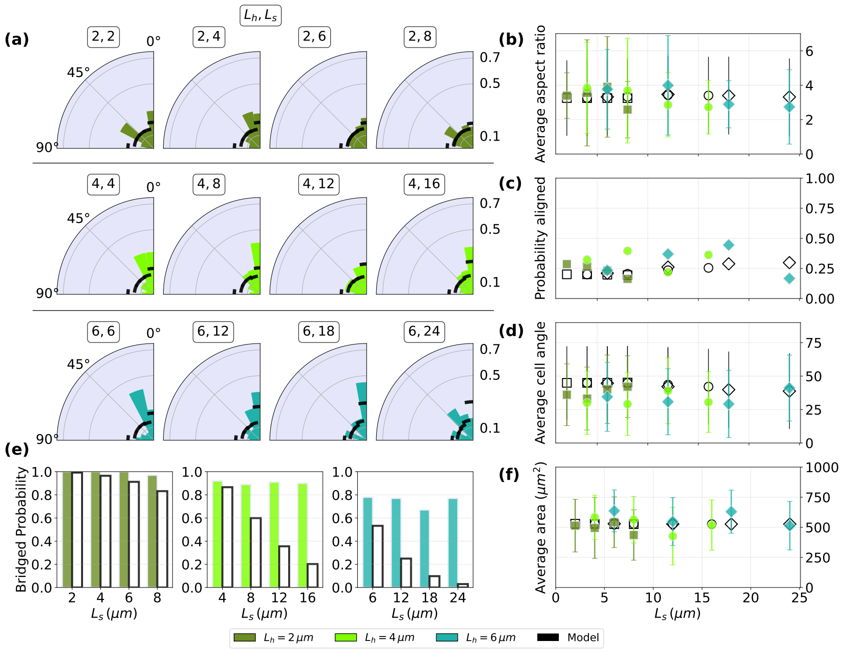

**Fig. SI-8** Contact guidance of HeLa cells deposited on patterns where soft stripes are softer then under standard conditions, to affect cell adhesivity. Comparison between experimental results (color coded by $L_h$) and model predictions (black, and hollow symbols) for varying widths of the soft substrate $L_s$. (a) Alignment of cells relative to the substrate stripes orientation . (b) Average cell elongation, reflecting changes in cell shape. (c) Percentage of aligned cells (d) Average cell orientation. (e) Probability for bridging between stiff lines. (f) Average cell area.

### 7 Tissues on stiffness patterns

#### 7.1 DPD simulations

We employ two-dimensional Dissipative Particle Dynamics (DPD) simulations with two-particle agents to investigate tissue growth [5]. Cellular dynamics is governed by various forces that account for both cell-cell and cell-environment interactions, while cellular membranes are reconstructed using Voronoi tessellations. This work extends the modeling framework previously introduced by the authors in [6, 7].

##### 7.1.1 Defining the stripes

We model alternating periodic stripes of hard and soft types, with widths $L_h$ and $L_s$, each characterized by a distinct dissipation coefficient. The stripe effect is incorporated via the intra-cell dissipative background force, which applies the appropriate friction coefficient in each region. Experiments are labeled as $L_h$-$L_s$.

To match simulations with experiments, a length of 5 simulation units is mapped to a stripe width of $4\mu m$ in the experiments, corresponding to the maximal size of edge cells in both systems.

##### 7.1.2 Initialization of the simulations

To avoid our results to depend on the initial positions of the cells, we use positions generated from a Poisson Disk Sampling. We performed the Poisson disk sampling using the SciPy implementation available in the `scipy.stats.qmc.PoissonDisk` class [8] based on Bridson's algorithm [9]. We selected the parameters of the algorithm so that the particles of the same cell do not overlap and stay in the distance range $[min\ D_{cp}, max\ D_{cp}]$. We also tune $min\ D_{cc}$ to avoid overlapping between particles of distinct cells. The range of initial velocity $min\ v_p$ to $max\ v_p$ is selected to prevent the same overlapping at the first time step. With the following set of parameters (Tab. 7.1.2) for the initial positions sampling, we are able to create a stable set of initial positions for each of our simulations.

| Name | Value | Description |
|---|---|---|
| $min\ D_{cc}$ | 1.0 | Minimum distance between cell centroids of two distinct cells during generation |
| $min\ D_{cp}$ | 0.2 | Minimum distance between individual cell particle and the centroid of the generated cell |
| $max\ D_{cp}$ | 0.4 | Maximum distance between individual cell particle and the centroid of the generated cell |
| $min\ v_p$ | 0.04 | Lower bound of randomly generated cell velocities |
| $max\ v_p$ | 0.1 | Upper bound of randomly generated cell velocities |
| $max\ t_0$ | 0.02 | To prevent synchronization of divisions at the very beginning, this controls how far into their initial state a cell can be initially |
| $min\ r_0$ | 0.8 | Minimum distance between particles at current state start |

##### 7.1.3 Parameters of the simulations

All model parameters are listed in the table below and the simulation snapshot are displayed in Fig. 6. Most parameters are recovered from previous work [6] where simulations were optimized to recover the tissue growth dynamics on gels. The only difference is in the parameter $R_t$ which is increased to improve the stability of the code, without quantitatively affecting the result.

| **Name** | **Value** | **Description** |
|---|---|---|
| $\mu_{bg}$ | 0.6 | Friction on the background/substrate |
| $\mu_{bg,h}$ | 0.6 | Friction on the hard stripe |
| $\mu_{bg,s}$ | 0.1 | Friction on the soft stripe |
| $L_h$ | *variable* | Width of the hard stripe |
| $L_s$ | *variable* | Width of the soft stripe |
| $B$ | 4.0 | Growth coefficient |
| $r_0$ | 1.12 | Constant offset for the calculation of the growth force |
| $\gamma_c$ | 1.1 | Dissipation within one cell |
| $R_t$ | 2.5 | Range of dissipative forces |
| $\Gamma$ | 0.0 | Active force coefficient |
| $k_a$ | 0.001 | Rate of apoptosis |
| $R_{c1}$ | 0.75 | Lower threshold for cell division |
| $R_{c2}$ | 10000 | Upper threshold for cell division |
| $r_c$ | 0.01 | Distance at which particles of a new daughter cell are placed after division |
| $k_B T$ | 0.424 | Thermal noise |
| $f_0$ | 150.0 | Repulsive cell-cell potential coefficient |
| $f_1$ | 1.0 | Attractive cell-cell potential coefficient |
| $R_{pp}$ | 1.0 | Range of pair potential coefficient |
| $\mu_{\parallel}$ | 0.2 | Dissipation coefficient between cells in parallel direction of connection-line |
| $\mu_{\perp}$ | 0.15 | Dissipation coefficient between cell perpendicular to connection-line |
| $m$ | $5.76 \times 10^{-4}$ | Cell particle mass for impulse and velocity calculation |
| $f$ | 1.4 | Factor in combination with $R_{pp}$, used for adhesion and neighbor search |
| $f_3$ | 1.0 | Density gradient force factor |
| $\tau_d$ | 0.25 | Time of division |
| $K_s$ | 1000.0 | Spring force coefficient to maintain cell size during division phase |
| $\lambda$ | $5 \times 10^{-6}$ | Lloyd optimization factor |
| $A_{max}$ | 18.34356 | Maximum area of the edge-cell polygon |

#### 7.2 MDCK monolayer spreading dynamics

##### 7.2.1 Experimental methods

For collective contact guidance, Madin–Darby Canine Kidney (MDCK) epithelial cell line, which are a gold-standard for epithelial model due to the ability to form highly polarized monolayers with mature tight junctions, recapitulate canonical apico-basal trafficking pathways, and collective migration behaviours closely resembling those of native epithelia. A clone that stably expresses LifeAct-EGFP was developed by the following: The construct (pLenti LifeAct-EGFP BlastR, Addgene #84383) was packaged into third-generation lentiviral particles and used to transduce subconfluent MDCK cultures. Forty-eight hours later, cells were placed under 5 $\mu$g mL$^{-1}$ blasticidin selection for seven days. Individual resistant colonies were cloned, expanded, and screened by epifluorescence microscopy; the brightest, most homogeneous clone was used for all subsequent experiments. Circular colonies 1.5 mm in diameter were seeded inside PDMS stencils on fibronectin-coated stiffness-stripe substrates, allowed to reach confluence for 18 h, and then released by gently lifting the stencil. Time-lapse imaging was performed in a 37 °C, 5 % $CO_2$ environmental chamber on a Nikon Ti-E inverted microscope fitted with a 10x objective. Phase-contrast and GFP fluorescence channels were captured every 1 h for 3 days.

##### 7.2.2 Experimental data treatment

Individual clusters were identified within each time frame using ImageJ analysis software. The images were first subjected to change to 8-bit, and background corrected to minimize the noise. Subsequently, the images were made binary, fitted with ellipse, and particle analyzer was used to measure the quantitative measurements of the morphological parameters, including individual cluster area, elongation, perimeter, and angle within different time frames. The analysis was conducted across various time frames for clusters grown on multiple strips (e.g. 2-2. 4-8).

Image analysis allowed us to extract the temporal evolution of the tissue's characteristics. In particular, we focus on the quantities of interest for comparison with our simulation results: the lengths of the major and minor axes of the tissue, and the angle between the major axis and the horizontal direction. Due to inherent noise in the experimental measurements of these quantities, we applied a Savitzky–Golay filter from scipy `scipy.signal.savgol_filter` class [8] to smooth the data. The filtered data are presented alongside the raw measurements for clarity. Below, we present the plots of these quantities (Fig.SI-9) .

#### 7.3 Analysis of simulations and experiments

##### 7.3.1 Total tissue area

Tissue area in the simulations is obtained from Voronoi diagrams of cell nuclei. To compare to the experiments, the timescale is fixed by matching area-doubling times in the early growth phase. Since simulated tissues are smaller than experimental ones due to computational limits, we compare relative area increases (Fig. SI-10), with

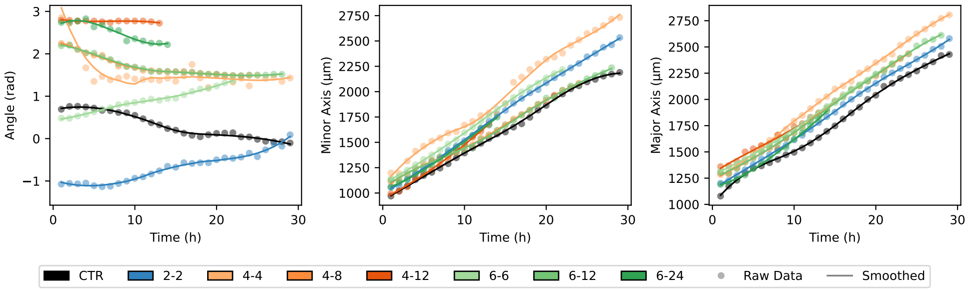

**Fig. SI-9** Multi-panel subplot displaying the experimental measurements used to compute the anisotropic growth parameter $\xi$. In each panel, raw data are shown as dots, while solid lines represent the smoothed data and their derivatives. The smoothed data were subsequently used to compute the corresponding derivatives, which in turn were employed to determine the order parameter $\xi$.

experimental data shown as points and simulations as lines. The left panel shows predominantly gel substrates, the right panel the glass control and mixed 50–50 substrates. As expected, predominantly gel substrates match better since parameters were tuned to mimic them [6]. A closer fit would require incorporating stiffness-dependent proliferation rates, which is beyond the scope of the current model.

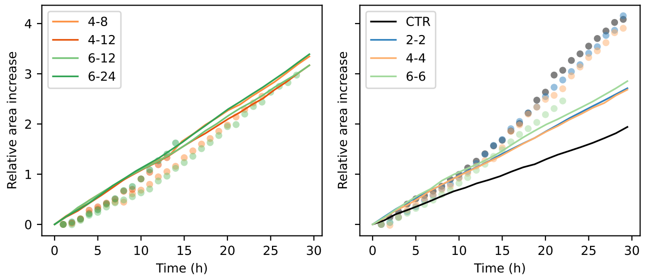

**Fig. SI-10** Relative change in tissue area, $(A(t) - A(t_0))/A(t_0)$, with the simulation timescale calibrated to experimental measurements. Simulation results are shown as lines, and experimental data as points. The left panel corresponds to predominantly gel substrates, while the right panel presents the glass control and mixed 50-50 substrates. Since the simulated proliferation rate was parameterized for the gel-like substrate as in [6], close agreement with experimental data is observed only in the left panel.

##### 7.3.2 Tissue elongation (major and minor axes)

To quantify tissue elongation, we computed the major and minor axes of the simulated tissue using the covariance ellipse of the cell nuclei coordinates, following the procedure in [10]. The normalized covariance ellipse has radii

$$r_x = \sqrt{1+p}, \quad r_y = \sqrt{1-p} \tag{S31}$$

where $p$ denotes the Pearson correlation coefficient[1]

The normalized ellipse is first rotated counterclockwise by 45°, and the normalization is reversed by rescaling the axes with the respective standard deviations multiplied by a precision factor $n$

$$s_x = n\sigma_x, \qquad s_y = n\sigma_y \ .$$

Finally, the ellipse is translated to the mean position, $(\mu_x, \mu_y)$. The factor $n$ determines the approximate confidence region captured by the ellipse[2], and here we use $n = 2.7$.

This procedure yields the axis lengths of the covariance ellipse but does not directly provide its orientation after rescaling. The derivation of the colony angle is described in the following section.

##### 7.3.3 Tissue orientation with respect to stripes

To account for the influence of stripes on the tissue, we also need the angle of the tissue relative to the stripes. We denote the desired angle as $\theta$, as shown in Fig. SI-11. To reproduce the procedure from the previous section, we start with the parametric equations of an ellipse with semi-axes $r_x$ and $r_y$, rotated by 45°

$$x(t) = \frac{1}{\sqrt{2}}(r_x \cos(t) - r_y \sin(t)), \quad y(t) = \frac{1}{\sqrt{2}}(r_x \cos(t) + r_y \sin(t)) \ . \tag{S32}$$

Next, we again use the scaling factors $s_x$ and $s_y$, which modify the equations to

$$x(t) = \frac{s_x}{\sqrt{2}}(r_x \cos(t) - r_y \sin(t)), \quad y(t) = \frac{s_y}{\sqrt{2}}(r_x \cos(t) + r_y \sin(t)) \ . \tag{S33}$$

Finally, to determine the angle $\theta$, we locate the vertices along the major axis by maximizing $x(t)^2 + y(t)^2$ using Mathematica [11] and obtain

$$R := r_x^2 + r_y^2 \quad S := s_x^2 + s_y^2 \tag{S34}$$

$$\tan(\theta + \pi/2) = \frac{-\left(R(s_x - s_y)(s_x + s_y) + \sqrt{r_x^4 S^2 + r_y^4 S^2 + 2r_x^2 r_y^2 (s_x^4 - 6s_x^2 s_y^2 + s_y^4)}\right)}{2(r_x - r_y)(r_x + r_y)s_x s_y} \ . \tag{S35}$$

##### 7.3.4 Growth velocities along and perpendicular to the stripes

In the experiments, the tissue is not necessarily aligned with the stripes and may change its orientation over time. Consequently, the elongation is no longer a reliable

[1]Pearson coefficient is defined as $p := \mathrm{Cov}[x, y]/(\sigma_x \sigma_y)$ where covariance is $\mathrm{Cov}[x, y] = \frac{1}{N}\sum_{i=1}^{N}(x_i - \mu_x)(y_i - \mu_y)$. A special case is the covariance of the same variable, known as variance $\mathrm{Cov}[x, x] = \sigma_x^2$.

[2]This is approximate because the region is elliptical rather than rectangular; see [10] for details.

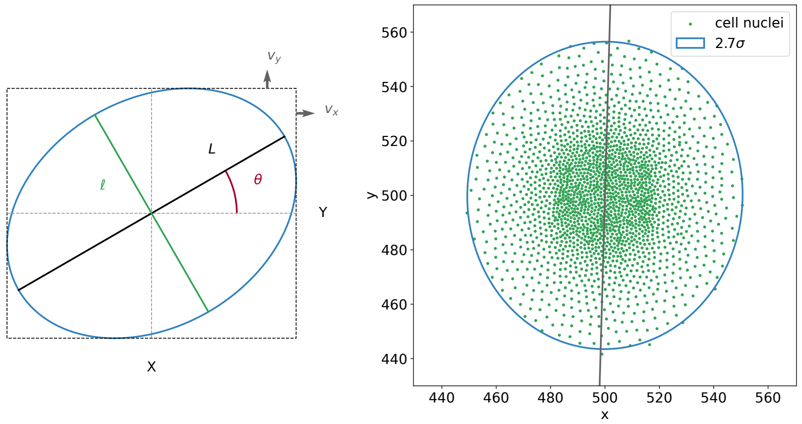

**Fig. SI-11** *Left:* Definition of parameters for a rotated ellipse centered at the origin and its bounding box. *Right:* Example of a confidence ellipse for the simulation data points with the major axis line.

measure of the influence of the stripes on the tissue growth. Instead, we analyze the change in tissue growth velocity both parallel and perpendicular to the stripes.
To quantify this, we define the bounding box of the ellipse with dimensions denoted as $X$ and $Y$, as illustrated in Fig. SI-11. For an ellipse centered at the origin with major and minor axes denoted by $L$ and $\ell$ respectively, and with a rotation angle $\theta$, the canonical equation is given by

$$\frac{(x\cos\theta - y\sin\theta)^2}{(L/2)^2} + \frac{(x\sin\theta + y\cos\theta)^2}{(l/2)^2} = 1 \; . \tag{S36}$$

To determine $X$ and $Y$, we differentiate $y$ with respect to $x$, yielding [12]

$$\frac{\partial y}{\partial x} = -\frac{L^2 x\sin^2\theta - y(L^2-\ell^2)\sin\theta\cos\theta + \ell^2 x\cos^2\theta}{L^2 x\cos^2\theta - x(L^2-\ell^2)\sin\theta\cos\theta + \ell^2 y\sin^2\theta} \; . \tag{S37}$$

The bounding box dimensions are obtained by setting the numerator and denominator to zero and solving these equations simultaneously with the ellipse equation. After algebraic manipulation, we derive the final expressions

$$X = \frac{1}{2}\sqrt{L^2\cos^2\theta + \ell^2\sin^2\theta}, \quad Y = \frac{1}{2}\sqrt{L^2\sin^2\theta + \ell^2\cos^2\theta} \; . \tag{S38}$$

Now we assume our ellipse evolves in time and all variables become functions of the time $t$ (i.e. we now have $X(t)$, $L(t)$, $\ell(t)$ and $\theta(t)$). Defining $A(t) \equiv L^2\cos^2\theta + \ell^2\sin^2\theta$ and differentiating using chain rule, we obtain

$$\dot{X}(t) = \frac{1}{4\sqrt{A(t)}}\,\dot{A}(t) \; . \tag{S39}$$

After further mathematical manipulation, we derive the expressions for the front velocities, $v_x = \dot{X}(t)$ and $v_y = \dot{Y}(t)$, used in the main paper

$$\dot{X}(t) = \frac{L\,\dot{L}\cos^2\theta + \ell\,\dot{\ell}\sin^2\theta + \cos\theta\,\sin\theta\,\dot{\theta}\,(\ell^2 - L^2)}{2\sqrt{L^2\cos^2\theta + \ell^2\sin^2\theta}}\,, \tag{S40}$$

$$\dot{Y}(t) = \frac{L\,\dot{L}\sin^2\theta + \ell\,\dot{\ell}\cos^2\theta + \sin\theta\,\cos\theta\,\dot{\theta}\,(L^2 - \ell^2)}{2\sqrt{L^2\sin^2\theta + \ell^2\cos^2\theta}}\,. \tag{S41}$$

###### 7.3.5 Growth anisotropy versus density profile

The goal of this analysis is to determine more precisely how stripe patterns affect the distribution of cell density and elongation within the tissue. In particular, we ask whether the observed elongation of the tissue arises primarily from an increased number of cells along the stripe direction, or whether it reflects elongation at the level of individual cells. To address this question, we analyze tissues simulated on a patterned substrate with $L_h = 12$ $\mu$m and $L_s = 6$ $\mu$m, using $\mu_{\mathrm{bg,h}} = 0.1$ and $\mu_{\mathrm{bg,s}} = 0.6$ (Fig. SI-12a). We focus on three quantities: (i) cell density, (ii) cell elongation, and (iii) cell orientation, measured both perpendicular to the stripes (X-axis) and parallel to the stripes (Y-axis). Each quantity is averaged within rectangular bins aligned with the two axes, and the results along the X-axis are shifted for comparison with those along the Y-axis, since the tissue extends less in the X-direction (Fig. SI-12b–d).

The analysis shows that the leading fronts of the density profiles are indistinguishable along the two axes (Fig. SI-12b). However, in the bulk, we see that locally cell morphology depends on substrate stiffness, causing nonuniform homeostatic density. This, combined with the geometry of the substrate, accumulates to produce the observed anisotropic elongation at the tissue scale.

The elongation and orientation distributions are also comparable along both axes, with only minor discrepancies (Fig. SI-12c,d).

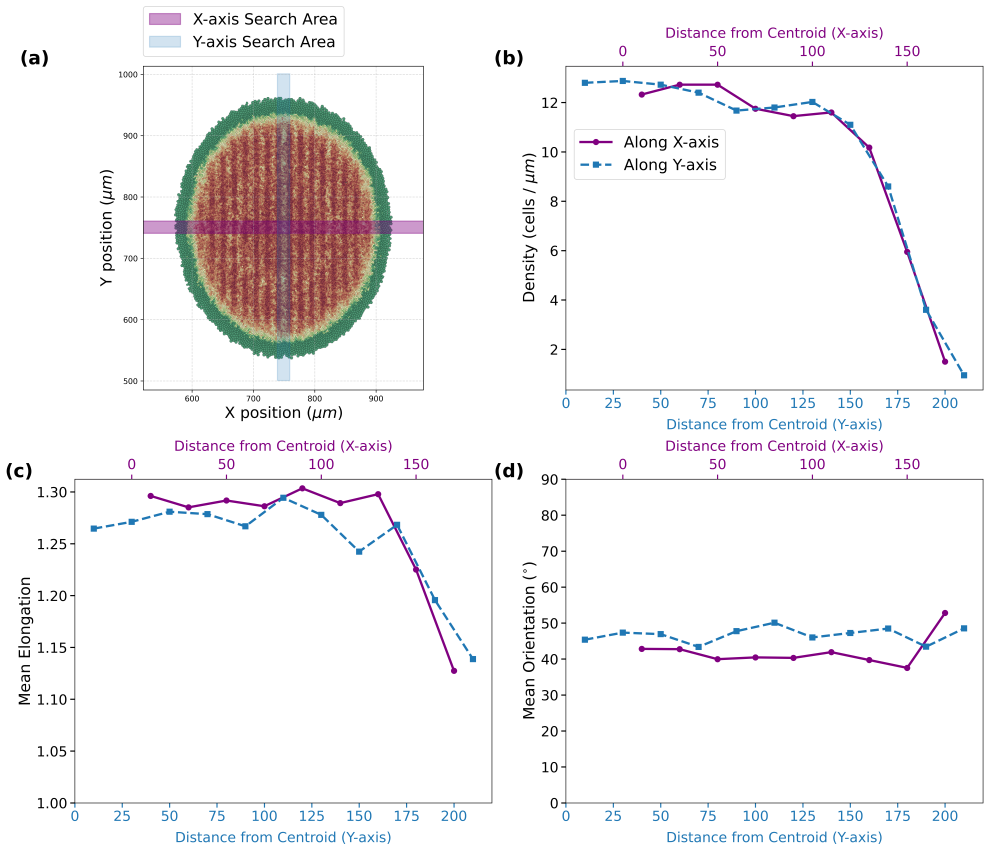

**Fig. SI-12** Analysis of tissue organization on a patterned substrate (simulation). (**a**) Voronoi tessellation of the simulated tissue. Color represents the cell size. Purple and blue shaded regions indicate the search windows used for computing properties along the $x$- and $y$-axes, respectively. (**b**) Cell density profiles along $x$ (purple) and $y$ (blue) directions. (**c**) Mean cell elongation as a function of distance from the centroid. (**d**) Mean cell orientation angle with respect to the $x$-axis.

[4] Avraham Moriel, Haguy Wolfenson, and Eran Bouchbinder. Characteristic energy

scales of active fluctuations in adherent cells. *Biophys. Rep. (N. Y.)*, 3(1):100099, 2023.

[5] Markus Basan, Jacques Prost, Jean-François Joanny, and Jens Elgeti. Dissipative particle dynamics simulations for biological tissues: rheology and competition. *Phys. Biol.*, 8(2):026014, 2011.

[6] Kevin Höllring, Lovro Nuić, Luka Rogić, Sara Kaliman, Simone Gehrer, Carina Wollnik, Florian Rehfeldt, Maxime Hubert, and Ana-Sunčana Smith. Capturing the mechanosensitivity of cell proliferation in models of epithelium. *Proceedings of the National Academy of Sciences*, 121(45):e2308126121, 2024.

[7] Sara Kaliman, Maxime Hubert, Carina Wollnik, Lovro Nuić, Damir Vurnek, Simone Gehrer, Jakov Lovrić, Diana Dudziak, Florian Rehfeldt, and Ana-Sunčana Smith. Mechanical regulation of epithelial tissue homeostasis. *Phys. Rev. X.*, 11(3):031023, 2021.

[8] Pauli Virtanen, Ralf Gommers, Travis E. Oliphant, Matt Haberland, Tyler Reddy, David Cournapeau, Evgeni Burovski, Pearu Peterson, Warren Weckesser, Jonathan Bright, Stéfan J. van der Walt, Matthew Brett, Joshua Wilson, K. Jarrod Millman, Nikolay Mayorov, Andrew R. J. Nelson, Eric Jones, Robert Kern, Eric Larson, C J Carey, İlhan Polat, Yu Feng, Eric W. Moore, Jake VanderPlas, Denis Laxalde, Josef Perktold, Robert Cimrman, Ian Henriksen, E. A. Quintero, Charles R. Harris, Anne M. Archibald, Antônio H. Ribeiro, Fabian Pedregosa, Paul van Mulbregt, and SciPy 1.0 Contributors. SciPy 1.0: Fundamental Algorithms for Scientific Computing in Python. *Nature Methods*, 17:261–272, 2020.

[9] Robert Bridson. Fast poisson disk sampling in arbitrary dimensions. In *ACM SIGGRAPH 2007 sketches*, page 22, 2007.

[10] Carsten Schelp. An alternative way to plot the covariance ellipse. https://carstenschelp.github.io/2018/09/14/Plot_Confidence_Ellipse_001.html, 2018. Accessed: 2025-02-13.

[11] Wolfram Research, Inc. Mathematica 11.0, 2016.

[12] Wenbo Dong, Pravakar Roy, Cheng Peng, and Volkan Isler. Ellipse R-CNN: learning to infer elliptical object from clustering and occlusion. *IEEE Transactions on Image Processing*, 30:2193–2206, 2021.
